# Spontaneously blinking fluorophores for accelerated MINFLUX nanoscopy

**DOI:** 10.1101/2022.08.29.505670

**Authors:** Michael Remmel, Lukas Scheiderer, Alexey N. Butkevich, Mariano L. Bossi, Stefan W. Hell

## Abstract

Spontaneously blinking fluorophores, a class of molecules switching rapidly between a dark and a brightly emitting state, have emerged as a popular core to build fluorescent markers for super-resolution microscopy. With typical on-times in the order of tens of milliseconds, they are most suitable for STORM and related nanoscopy methods. Recent MINFLUX nanoscopy, however, can localize molecules even within a millisecond and achieve an up to ten times higher localization precision. Here, we present a series of spontaneous blinkers with short on-times (1-3 ms) matching MINFLUX recording time-scales. Our design builds upon a silicon rhodamine fluorescent core with a modified thiophene- or a benzothiophene-fused spirolactam fragment, which shifts the spirocyclization equilibrium toward the dark closed form at physiological conditions, imparting cell permeability. Concurrently, we obtain a highly photostable, short-lived open form with bright red emission. Characterizing the blinking behavior of single fluorophores bound to three different protein tags (antibodies, nanobodies, and HaloTag self-labeling enzyme) allowed us to select the best candidate for MINFLUX microscopy. The short on-times speed up MINFLUX localization by up to 30-fold.

---

Super-resolution fluorescence microscopy (nanoscopy) afforded the all-optical visualization of specimen details much finer than the ~250 nm diffraction resolution limit.^1^ The first generation of these techniques (STED, RESOLFT, PALM, STORM, PAINT)^2, 3, 4, 5, 6, 7^ can routinely achieve a resolution down to 20-50 nm, enabling the detailed visualization of submicrometer-size cellular structures in (living) cells.^8^ While they all use two distinct fluorophore states for telling neighboring fluorophores apart (typically a fluorescent ‘on’ and a dark ‘off’ state), nanoscopy methods differ in the spatio-temporal control of the ‘on’ and ‘off’ states and the way the position of the fluorophore is established in the sample. The coordinate-stochastic single-molecule methods called PALM, STORM, and PAINT infer these positions from the diffraction pattern of the fluorescence light rendered by single fluorophores on a pixilated detector such as a camera. In contrast, the coordinate-targeted methods STED and RESOLFT scan the sample with a spatially modulated illumination pattern with a minimum intensity node (ideally zero) that defines a reference position for the markers. In single beam scanning implementations this pattern is typically shaped like a donut. By synergistically using the unique advantages of both strategies, the recently introduced MINFLUX (MINimal photon FLUXes) concept pushes the fluorescence nanoscopy resolution down to a few nanometers. Concretely, MINFLUX separates neighboring fluorophores by on/off-switching their emission capability individually, like PALM or STORM, but localizes the emitting fluorophore with a pattern of excitation light featuring an intensity node, like the donut used in STED microscopy. Localization with a donut-shaped excitation beam shifts the burden of requiring many photons for localization on the excitation beam, so that the number of fluorescence photons required for attaining a certain localization precision or speed is substantially reduced. Minimizing the required number of detected photons lessens the demand on the on/off-switchable fluorophores regarding brightness in the ‘on’-state and photostability. Thus, the possible range of viable fluorescent molecular switches and switching mechanisms is expanded over that used in established PALM and STORM techniques.

In principle, the fluorophores that work for PALM and STORM, such as cyanine dyes,^9, 10^ or caged rhodamines^11^ also work for MINFLUX microscopy,^12, 13, 14, 15, 16^. Yet, as they are optimized for the detection of isolated fluorescence patterns on a camera, typical PALM/STORM fluorophores do not necessarily leverage the specific advantages provided by MINFLUX in terms of attainable resolution and speed. The on-time (T_ON_) preferably matches the frame-rate of the camera (T_ON_ ≈ frame-rate^-1^ ≈ 10-20 ms), and the fluorophore has to provide a large number of detected photons (typically N_PH_ = 1000-5000) during the on-time. In contrast, MINFLUX requires less than a tenth of that number of photons to achieve the same resolution^12^ and can localize emitters in time intervals below a millisecond.^17^ Besides, cyanine and caged triarylmethane fluorophores entail unfavorable conditions. For example, the blinking behavior of cyanines requires complex redox buffers, comprising an enzymatic system and several chemicals (thiols or reducing agents).^18^ Thus they are incompatible with live-cell imaging and their imaging conditions may affect other markers or functional biomolecules. It is widely recognized that the imaging buffer affects the image quality. Therefore, a wide variety of buffer compositions^19,20,21,22,23^ have been used; some of them are even undisclosed. For multicolor applications, pairing a cyanine (reporter fluorophore) with a suitable auxiliary dye (activator) was also reported.^24^ Besides rapid photobleaching, cyanines are prone to oxygen-dependent chemical photobluing stemming from the shortening of their polymethine chain by two carbon atoms^25^. This adverse feature challenges their application for multicolor imaging.^26^ Despite their successful use in super-resolution techniques for more than a decade, the physicochemical mechanism of the blinking behavior is still debated.^18, 27, 28^ While caged fluorophores have simpler, more reliable, and usually known activation mechanisms (on-switching)^29, 30, 31, 32, 33, 34, 35^, their off-switching relies on complex and poorly defined photobleaching reactions. Moreover, their activation can be induced by the excitation laser (in a one or two-photon process), and the photoactivation products (rhodamine dyes) are prone to oxidative photobluing.^36^ Finally, the reaction kinetics of both on- and off-switching of cyanines and caged fluorophores strongly depend on the irradiation wavelength and intensity. In view of this complexity, fluorophores whose blinking is induced solely by thermal reactions (and hence involving only ground states) become an important alternative. While the use of such spontaneous blinking or self-blinking dyes is well established in STORM and related methods^37, 38, 39, 40, 41, 42, 43, 44, 45, 46, 47, 48, 49^, their exploitation in MINFLUX^50^ is still uncharted.

Here, we report on a series of spontaneously blinking fluorescent labels based on a reversible spirocyclization of silicon rhodamine amides. We systematically study their physicochemical properties, with emphasis on their blinking behavior on the single-molecule level. Emphasis is placed on the characteristics that are relevant for MINFLUX nanoscopy. To exploit the high localization speed and the rather low number of fluorescence photons required in MINFLUX, we looked for compounds with short T_ON_ (< 10 ms) and moderate N_PH_ rather than for high duty cycles and fluorescence photon yields. In addition, we assessed the influence of the blinking on the T_OFF_ and the duty cycle (DC = T_ON_(/[T_ON_+T_OFF_]), as the blinking process imposes a limit on the maximum number of distinguishable markers in a diffraction-limited light spot. As a fluorescence microscope images just fluorophores but not the biomolecules, this so-called maximum ‘labeling density’ limits the translation of the localization precision into the separation of labeled biomolecules at the nanoscale.^51^ We observed certain variations of the relevant parameters by the nature of the labeled biomolecule (protein) used for targeting desired structures. While this variability is consistent with the effects of the local environment on the position of the spirocyclization equilibrium, it emphasizes the importance of the selection of appropriate tagging or labeling methods. We prepared reactive adducts for their application for fixed- and live-cell common labeling strategies, with conventional methods using bioconjugates (nanobodies) and a self-labeling enzyme (HaloTag). Finally, with the best identified candidate we imaged nuclear pore complexes (NUPs) with a fluorophore localization precision <3 nm using a commercially available MINFLUX microscope. We paid particular attention to the attainable localization efficiency and speed, analyzing the effect of blinking properties on the imaging.

## RESULTS and DISCUSSION

### Design and synthesis of spontaneously blinking fluorophores

The majority of reported spontaneously-blinking fluorescent labels are derived from a rhodamine core. Rhodamine dyes afford brightness and photostability, a wide spectral range (green to near-IR emission, with excitation from 488 nm to 640 nm) and a relatively simple and well-documented synthetic chemistry. While they are also capable of intramolecular spirocyclization between their electrophilic xanthylium chromophore and the 2’-carboxylate nucleophile to form a colorless and non-emissive lactone, in polar environments the equilibrium is highly shifted toward the emissive zwitterionic form. Bearing a stronger 2’-carboxamide nucleophile, the corresponding rhodamine amides exist nearly exclusively in the form of colorless spirolactams. As the thermal blinking of these early rhodamine spirolactams is inefficient at physiological pH, STORM has been carried out with these dyes by photoactivation.^52,53^ Custom chemical design yielded spontaneously blinking derivatives of 2’-(hydroxymethyl)silicorosamine (HMSiR)^38^, sparing 355-405 nm UV photoactivation. In further reports, both expansion of the spectral range^54^ and rational tuning of their blinking behavior^55^ were reported for this class of fluorophores.

In our design, we started from the silicon rhodamine (SiR) core structure, because the excitation and emission of SiR dyes in the red spectral region matches the 640 nm laser line of commercial microscopes. Red light also minimizes background in cell imaging, keeps photodamage and light dispersion at bay, and increases cell and tissue penetration. However, the amido derivatives of the SiR dye itself (Figure 1A, Ar_1_ = 1,2-phenylene) do not undergo spirolactam ring opening in aqueous buffers. To shift this equilibrium toward the open form and tune the blinking behavior, we explored a series of SiR-derived amides with varying Ar_1_ and Ar_2_, following the strategy that has been successfully used to optimize the fluorogenicity and cell membrane permeability of a series of rhodamine derivatives.^47, 53, 56, 57^ Replacement of a fused benzene ring in the spirolactam core (Ar_1_) with a thiophene or a benzothiophene resulted in the desired spontaneous blinking behavior with a far-red (680-690 nm) fluorescent emission (Figure S1). Further tuning of the electronic properties of the *N*-amido substituent (Ar_2_) allowed the desired control over *K*_a2_ equilibrium constant (Figure 1B)^58, 59^ and, as a result, over the extent of spirolactam ring opening in neutral aqueous buffer and the characteristics of the stochastic blinking. Compounds **1-6** were prepared from the corresponding SiR lactones with modified fused aromatic ring, which were converted to the corresponding acyl chlorides upon treatment with oxalyl chloride or phosphorus(V) oxychloride and condensed with the appropriately substituted 3- or 4-aminobenzoic acid ester (7-aminocoumarin derivative for **5**). Free carboxylic acid forms of the dyes **1-6** were recovered by mild acidic or basic hydrolysis (see Supporting Information for the experimental details). Further functionalization of the carboxylic acid provided the amino-reactive *N*-hydroxysuccinimide (NHS) esters, thiol-reactive maleimide derivatives and self-labeling HaloTag protein (an engineered ω-chloroalkyl dehalogenase^60^) ligands.

**Figure 1.**
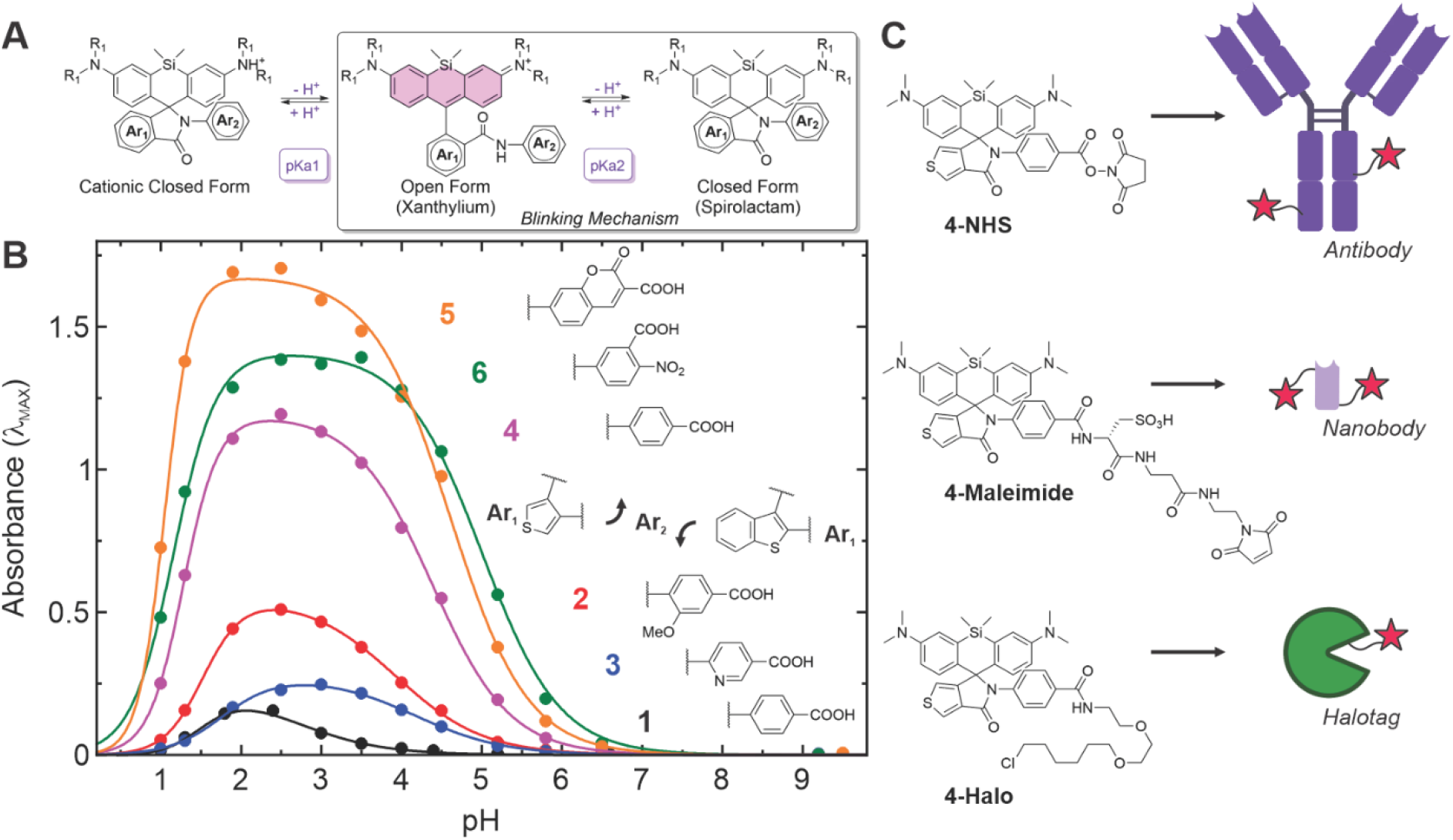
(A) General structure of the studied fluorophores and the reactions responsible for their blinking behavior; (B) pH dependence of the red absorption band of the closed form and the structures of the substituents Ar_1_ and Ar_2_ for each compound (dots: measurements; lines: fits); (C) Reactive adducts (NHS, maleimide, chloroalkane) and schematic representation of the labeling strategies (antibodies, nanobodies and self-labeling enzyme HaloTag).

**Figure 2.**
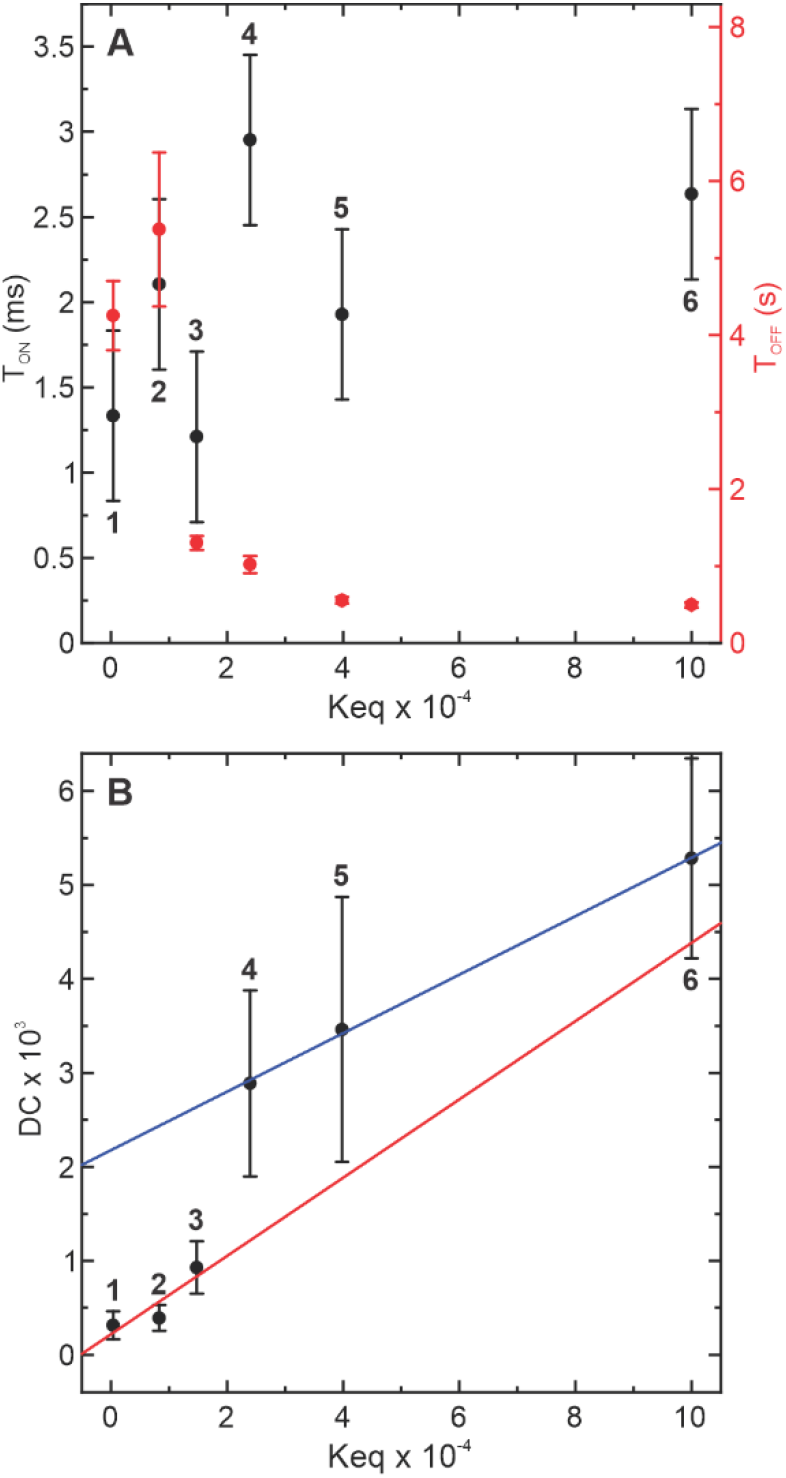
(A) Average on-times (T_ON_), off-times (T_OFF_), (B) and the resulting duty cycle (DC) measured from single molecule traces of antibodies labeled with compounds **1-6** with a low degree of labeling (DOL ≤0.3), as a function of the equilibrium constant (K_eq_ = 1/K_a1_). A linear regression is plotted for each group of compounds (**1-3**: Ar_1_ = benzothiophene, **4-6**: Ar_1_ = thiophene).

### Ensemble characterization of carboxylic acids 1-6 in solution

We first studied the absorption and emission properties of the six compounds in a buffered solution at pH = 7.4 (Figure S1). The effect of Ar_2_ substituent pattern on the spectral properties was minimal, with maximum absorption and emission variability within 10 nm. The emission efficiency also shows small changes (Table 1), with benzothiophene derivatives demonstrating decreased fluorophore brightness. On the contrary, considerable variations were observed in response of the spirolactamxanthylium amide equilibrium to the change of pH (Figure 1 and S2), with the spirolactam form predominant in neutral and basic environments, and the xanthylium in moderately acidic media (corresponding to p*K*_a2_, see Figure 1A and Table 1). This equilibrium is more shifted towards the spirolactam closed form for Ar_1_ = thiophene than for Ar_1_ = benzothiophene. Within each group, varying the electronic properties of Ar_2_ results in the fine-tuning of the equilibrium. Below pH = 2, a second colorless and non-emissive form becomes prevalent for all compounds, suggesting a second protonation step corresponding to p*K*_a1_. While this behavior was observed for some rhodamine amides^53^ and HMSiR analogues,^40, 61^ it has not been described for SiR-amide derivatives.^47, 57^ The proposed species and involved equilibria are shown in Figure 1A. Based on this simplified representation, the two equilibrium constants were extracted by fitting the absorbance at the maximum of the xanthylium form, with a Hill’s model ^61, 62^. It is noteworthy that all compounds predominantly exist in the spirocyclic closed form (SP, Figure 1A) at physiological pH values. This neutral form is known to be cell permeable for similar rhodamines and silicon rhodamines.

**Table 1.** Photophysical properties of compounds 1-6.

| Comp. | $\Phi_{\text{Fluo}}$ | $n_1$ | $pK_{a1}$ | $n_2$ | $pK_{a2}$ | $K_{eq}$ | DC<br>X1000 | $T_{ON}$<br>(ms) | $T_{OFF}$<br>(s) | $N_{CY}$ | %BI | $PH_{CY}$ | Rate<br>(kHz) |
| --- | --- | --- | --- | --- | --- | --- | --- | --- | --- | --- | --- | --- | --- |
| 1 | 0.10 | 1.70 | 1.62 | 0.79* | 2.62 | 417 | 0.31 | 1.3 | 4.25 | 13 | 45 | 143 | 111 |
| 2 | 0.17 | 1.52 | 1.52 |  | 3.92 | 8318 | 0.39 | 2.1 | 5.37 | 7 | 62 | 251 | 120 |
| 3 | 0.24 | 1.77 | 1.77 |  | 4.17 | 14791 | 0.93 | 1.2 | 1.30 | 36 | 53 | 115 | 96 |
| 4 | 0.16 | 1.28 | 1.28 | 0.90* | 4.38 | 23988 | 2.89 | 3.0 | 1.02 | 14 | 86 | 354 | 118 |
| 5 | 0.17 | 1.05 | 1.05 |  | 4.60 | 39811 | 3.46 | 1.9 | 0.56 | 39 | 78 | 207 | 109 |
| 6 | 0.16 | 1.28 | 1.28 |  | 5.00 | 100000 | 5.28 | 2.6 | 0.50 | 58 | 71 | 278 | 107 |

$K_{eq} = (K_{a1})^{-1}$ ; $n_{1,2}$ : Hill coefficients (\* global fit); $T_{ON}$ (average time, from bi-exponential fits); $T_{OFF}$ (from mono-exponential fits); $DC = T_{ON} \cdot (T_{ON} + T_{OFF})^{-1}$ , $N_{CY}$ : cycles measured in 100 s (WF), %BI = $100 \times (N_{CY} / N_{CY}^{TH})$ ; $PH_{CY}$ (Confocal); rate = $PH_{CY} / T_{ON}$ (Confocal).

### Single molecule characterization

The presence of two separate prototropic equilibria and the proximity of the values *K*_a1_ and *K*_a2_ makes it difficult to corroborate if there exists a pH interval within which any of the SiR-amides **1-6** is present solely in the xanthylium form. For this reason, the molar absorption coefficients of the colored form could not be reliably calculated. Moreover, the absorption of dyes **1-6** is too low and close to the instrument noise levels at neutral pH, so the duty cycles below 10^−3^ are expected. It was therefore deemed practical to evaluate these parameters by studying the blinking properties at the single molecule level. First, full IgG secondary antibodies were treated with reactive NHS ester derivatives of **1-6** (Figure 1C) to a low degree of labeling (≤0.3) to minimize the probability of tagging a single protein with more than one blinker. The antibodies were affixed to a cover slide based on a streptavidin/biotin double recognition system (Figure S3), whose extremely high affinity prevents any detachment of the antibodies for even longer times than those required for the experiment. Concentrations and incubation times were optimized to ensure a sparse surface distribution of the antibodies labeled with blinking fluorophores. Finally, samples were mounted in PBS and imaged on a custom-made microscope allowing for widefield (camera-based) and confocal detection. With the first, parallel data collection (ca. 30 molecules in a 12 × 12 µm area) was achieved with 10 ms temporal resolution, and with the second one a higher time-resolution of 1 ms was reached. Excitation intensity was the same in both modalities, and approximately matched the values necessary for imaging in a MINFLUX microscope (66 kW/cm^2^, calculated in the sample’s plane). Due to the large difference in the characteristic times of the two processes (on- and off-switching), we extracted T_OFF_ from wide-field experiments and T_ON_ from confocal measurements. However, the analysis procedure for the data from single molecule traces was the same in both cases (Figure S4-S7). The data is summarized in Table 1 and Table S1, along with other relevant calculated parameters. Since not all molecules are bleached at the end of the experiment, calculated N_C_ should be considered a lower limit.

In general, benzothiophene-derived spirolactams **1-3** demonstrate shorter on-times and longer off-times (and therefore a lower duty cycle) than thiophene derivatives **4-6**. While T_ON_ and T_OFF_ under neutral conditions do not have a clear correlation with the equilibrium constant *K*_a2_ corresponding to the reversible spirocyclization, a reasonable correlation was observed for the DC within each group. The T_OFF_ parameter demonstrates larger variation than T_ON_ and thus has a stronger effect on DC. Compared to other blinking fluorophores used in super-resolution techniques (including cyanines),^19^ the DC of compounds **1-6** lies in the same range of 10^−3^ – 10^−4^. Such low values are known to be compatible with stochastic methods^6, 19^ and MINFLUX^12, 15^ imaging with high labeling density. In addition, we found no observable changes in T_OFF_ upon illuminating the sample with UV light of 405 nm with intensities up to 1kW/cm^2^. We therefore dismiss a potential contribution of a photo-induced activation/on-switching mechanism under our conditions (aqueous environment, pH = 7.4, irradiation wavelength and power). With respect to the DC, there is little information in the literature of spontaneously blinking fluorophores to compare.^37, 38, 40, 42, 43, 44, 47, 50^ Regarding on-times, compounds **1-6** present DC values that are in general 1 to 2 orders of magnitude shorter than for previously reported compounds (Figure S6).^38, 39, 43, 47^ They also present non-mono-exponential behavior: after fitting to a bi-exponential model, we found a predominant short T_ON_ component at the limit of the experimental temporal resolution (0.5 – 1 ms), and the long component with T_ON_ ≈ 3 – 10 ms. These are slightly shorter than the typical localization times used in MINFLUX microscopy,^14, 15^ and considered too short for purely stochastic methods. Nevertheless, we obtained STORM images with fixed and live cells (Figure S8-S9) using immunostaining and HaloTag labeling, respectively. The latter also confirms the cell-permeability of HaloTag ligands derived from compounds **3-6**.

For MINFLUX imaging, we initially selected thiophene derivatives **4-6** for their shorter DC (ca. one order of magnitude shorter than benzothiophenes **1-3**). In this group, we selected the compound **4** with a slightly longer T_ON_ than the others. Next, three distinct tags, commonly employed for labeling biological samples, were selected (Figure 1C): NHS ester for tagging free primary amino groups of secondary antibodies (for imaging purposes, higher DOLs ≈2-3 were prepared), maleimide for selective modification of cysteine residues in engineered nanobodies, and linear ω-chloroalkane with a short PEG2 linker for labeling HaloTag fusion proteins. In the case of nanobodies, we have noticed that several hours post-labeling with the hydrophobic dyes, significant aggregation and precipitation of the tagged protein from the aqueous solution makes their application impractical. This solubility problem was solved by the introduction of an additional hydrophilic dipeptide linker (Cya-β-Ala, termed a “universal hydrophilizer”^63^), obtaining complete labeling to a defined DOL of 2 (Figure S10A). To confirm the covalent binding of a HaloTag ligand to its target fusion protein, ESI mass spectroscopy of a labeled HaloTag7 sample was performed (Figure S10B). The blinking behavior of fluorophores in these biomolecular complexes was also studied (Figure S7, Table S1) and compared with the correspondingly labeled antibody. We also confirmed their suitability for STORM imaging (Figure S11). We observed a considerable influence of the environment on the on- and off-times and therefore on the resulting duty cycle, especially for the HaloTag conjugate. To our knowledge, there is not much information in the literature regarding the effects of labeling on the duty cycle of blinking fluorophores, probably because of a marginal effect on imaging with a camera detector, given the long exposure times (typically >10 ms). However, we expect a larger impact on MINFLUX imaging, in particular when applying much shorter localization times (≤ 10 ms). Intrigued by the non-mono-exponential behavior, we then decided to inspect the on-times in detail. While a bi-exponential model fits the data and allows calculation of average on-times, a bimodal distribution may not be an adequate physical description of the system, with rather complex microenvironments affecting the fluorophore. We found a better comparison by a model-independent analysis of the experimental complementary cumulative distribution function (CCDF) of T_ON_ (Figure 3). In the first place, we observed an initial drop of 50-65% that accounts for the fraction of T_ON_ ≤ 1ms. These are probably discarded in camera-based localization methods (because of low photon numbers or STB level), but can be potentially localized in MINFLUX; this will be discussed in the next section. Another large group of T_ON_’s (25%-40%) is up to ~10 ms for labeled HaloTag and nanobodies or ~20 ms for antibodies, which corroborates the suitability of compounds **1-6** for STORM imaging.

**Figure 3.**
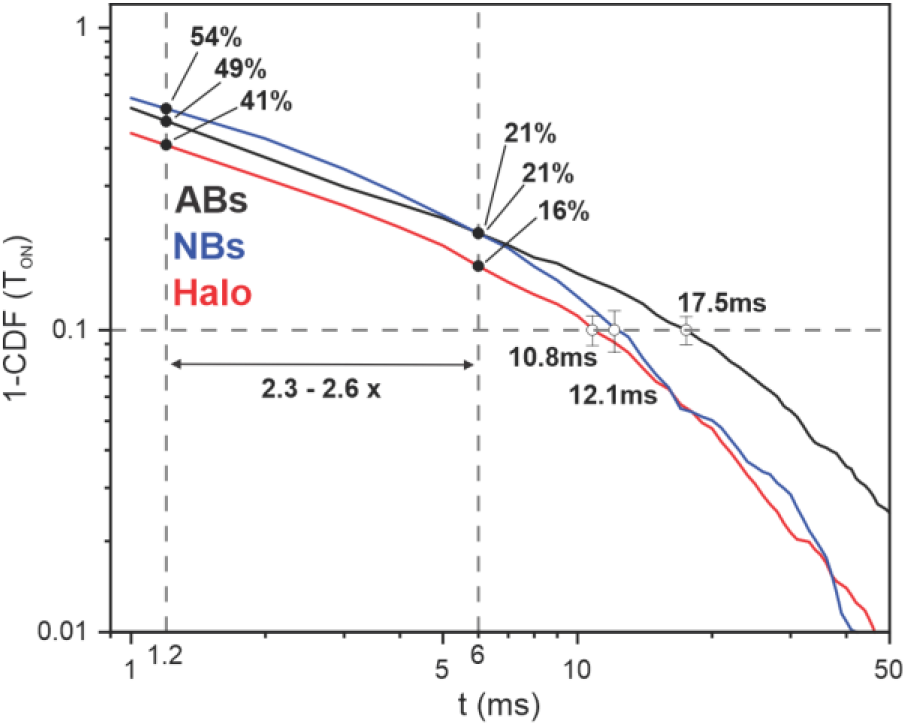
Complementary cumulative distribution function of T_ON_ for compound **4** measured in antibodies (black), nanobodies (blue) and bound to HaloTag7 (red). The values at 1.2 ms and 6 ms (see imaging sequences), and the crossing points at 10% are marked for each case.

### MINFLUX Imaging

To show their suitability for MINFLUX imaging, we labeled mEGFP-Nup107 cells with **4** linked to an anti-GFP nanobody through a hydrophilic linker and U2OS-Nup96-Halo cells with **4-Halo** ligand. Nuclear pore complexes (NUPs) have been previously proposed and used as a benchmark for super-resolution^15, 64^. The imaging was performed on a commercial MINFLUX setup without any instrumental modifications (see Supplementary Materials for details). Initially, a standard MINFLUX sequence optimized for cyanine dyes (e.g. Alexa Fluor 647, Sulfo-Cy5, Biotium CF660 and CF680) and provided by the manufacturer was used. The localization process can be split in two parts (see Table S2 for details). The first part consists of a series of 4 steps to pre-localize a molecule and progressively lower the MINFLUX localization range *L*,^12, *14*^; *it lasts 4 ms and requires at least a total of 120 photons above background. The second one consists of a pre-localization step (L* = 76nm) and the final localization step (*L* = 40 nm), rendering the position of the molecule. The dwell time is 2 ms (1+1 ms) and requires at least 30 photons per step. This second part is repeated as long as the molecule is in the on-state. Thus, each single molecule event (trace) results in a series of N_LOC_ localizations and a total number of photons N_PH_ over a localization time T_LOC_. The localization precision is evaluated from the dispersion of the localization of all molecules around their corresponding mean position.^14, 15, 16^ For both staining strategies, with anti-GFP nanobodies and HaloTag, we obtained MINFLUX images with compound **4**, comparable to those previously reported with a similar setup (Figure 4).^15^ Without localization binning, a localization precision of *σ* = 2.6 *nm*/*σ* = 2.9 *nm* was reached (Figure 4 B and D). From further analysis of the traces, we estimated for each SM trace a median of 7/8 localizations, a total of 1300/1300 photons (at a rate of 57kHz/45kHz) per molecule trace (185/163 photons per localization) for the nanobody/HaloTag respectively (Table S3). Previous work reported a localization precision below 1 nm calculated from traces with at least 4/5 localizations with 2100/2000 photons/loc^14, 15^. At an emission rate of 30/50 kHz, a molecule should persist in the on-state for at least 260/200 ms to be localized with such precision, corresponding to an average operative T_LOC_ (i.e. for blinking events effectively used to achieve such localization precision) of several hundreds of milliseconds (Table S3). Since the on-time of cyanines is typically several tens to a few hundreds of milliseconds (power dependent),^27, 28^ a large fraction of blinking events will be rejected with a standard imaging sequence, because they do not produce the required number of photons. Compounds **1-6** have on-times in the range of few milliseconds, leaving an even smaller fraction of fluorophores localizable in the on-state for more than 10-20 milliseconds (Figure 3). From the distribution of the measured on-times (Figure 3 and Figure S7), we conclude that images acquired with this standard (“slow”) routine (Figure 4) result predominantly from the fraction of blinking events with the long T_ON_. To evaluate the obtained localization precision, we binned the data presented in Figure 4, to compare with the precision values reported previously (Figure S12). Binning with a set value of 350 photons, we reached a raw localization precision of 1.9 nm, which can be reduced to 1.1 nm after applying a filtering method (DBSCAN + outlier suppression, for details see Materials and Methods section). However, binning the data acquired with our blinkers results in a loss of events (rejected due to a short T_ON_ and thus low N_PH_), affecting the quality of the image.

**Figure 4.**
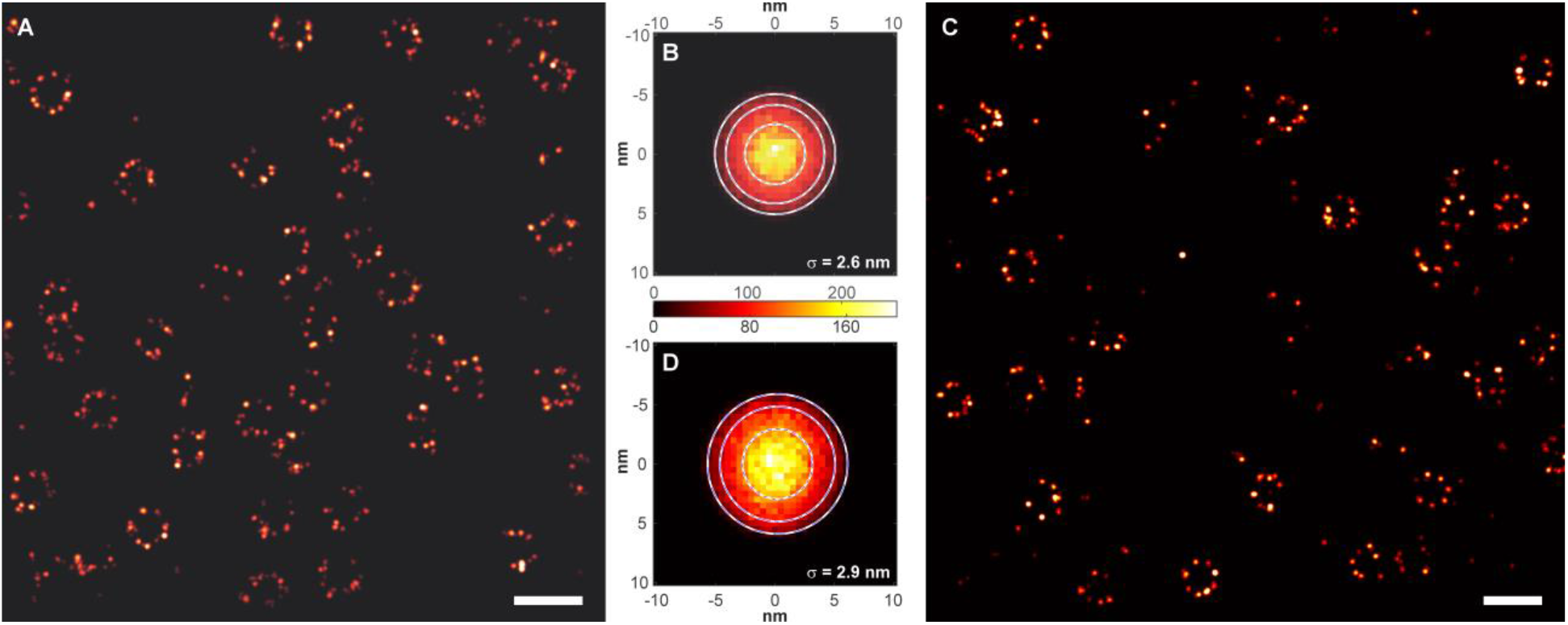
MINFLUX images of (A) mEGFP-Nup107 fixed cells labeled with an anti-GFP nanobody (dye **4**, DOL = 2), and (C) U2OS-Nup96-Halo fixed cells labeled with **4-Halo** (live-cell labeling and then fixed, see Materials and Methods section for details). The corresponding 2D dispersion plots (distance between localization and mean position of the cluster) are presented in (B) and (D), respectively. Scale bar: 200 nm.

These initial results showed that the spontaneously blinking fluorophore **4** could be successfully used in MINFLUX imaging and required a total acquisition time similar to previously reported values for cyanine labels. However, they did not take advantage of the full potential of our dyes as fast blinkers. In fact, setting a high bar on the total photon budget (up to 8000) is at the expense of localization speed (up to several hundreds of milliseconds per localization, on average) and bears the risk of fluorophore dislocation. The precision required for discerning single biomolecules (ca. 2-4 nm, the size of many proteins such as GPF, HaloTag or a nanobody) can be achieved with much fewer photons (< 100) in MINFLUX.^12^ Keeping an eye on localization speed, we decided to adapt the MINFLUX sequence to the photophysical properties of our fast blinkers (in particular their short on-times) and make use of a larger fraction of blinking events with short on-times. The main optimizations of the MINFLUX imaging routine were: (1) lowering the pre-localization time from 4 ms to 0.7 ms by reducing the number of steps from 4 to 2 and the localization time from 2 ms to 0.5 ms, and (2) increasing the localization range *L* from 40 nm to 76 nm for the localization step. With these modifications, the minimum time required to localize a molecule was reduced from 6 ms to 1.2 ms. To compensate for the loss of photons, the excitation intensity was increased by a factor of 2, to a value close to the maximum of the laser power available in the commercial microscope. Overall, we expected a trade-off in spatial resolution (particularly due to a 2.5× increase in *L*) for a 5-fold increase in localization speed and the fraction of blinking events used.

An image acquired with the optimized sequence (“fast”), directly compared with the “slow” sequence image (both at the same excitation power) is presented in Figure 5 (a temporal image buildup is shown in Figure S13), for mEGFP-Nup107 labeled with anti-GFP nanobodies labeled with dye **4**. The emission rate of the fluorophores was enhanced to 76 kHz and 118 kHz for the slow and the fast sequence, respectively. An average value of 314 and 75 photons/localization was obtained, resulting in a localization precision of 2.3 and 3.7 nm, respectively (Table S3). This moderate loss of resolution (by a factor of 1.6) is accompanied with a 5-7-fold increase in the effective average localization speed (Table S3, T_LOC_) and improved localization efficiency. The latter can be concluded from the total events on the images on Figure 5, normalized by the numbers of NUPs, showing a ≈2.6-fold increase for the fast sequence. Moreover, the cumulative distribution function shows that 90% of the traces were acquired with the fast sequence within 42 ms (on T_LOC_ distribution), closer to the 12 ms (on T_ON_ distribution) obtained from single molecule characterizations (compare Figure 5 and Figure 3). We attribute this difference to three distinct effects. First, at least a minimum of 2 localizations per molecule, plus the pre-localization, are necessary (see SI methods), limiting our time resolution to 1.7 ms as compared to 1 ms in the single molecule experiments. Second, despite approximate matching of the total excitation intensity between experiments, the fluorescent molecule experiences a variable effective value (probably lower) as it is moved near the nodal point in the imaging experiment. Longer blinking events producing larger detection numbers are favored, pushing the observed T_LOC_ to larger values. Finally, despite our best efforts to maintain the same molecular micro-environment in the characterization and imaging experiments, small differences cannot be excluded. Compared to previous work based on cyanine fluorescent labels, we increased the speed of localizations by a factor of 20-30× with only a 2-4-fold decrease in localization precision. Nevertheless, the final localization precision of our fast (optimized) sequence is still below the typical size of the protein tags.

**Figure 5.**
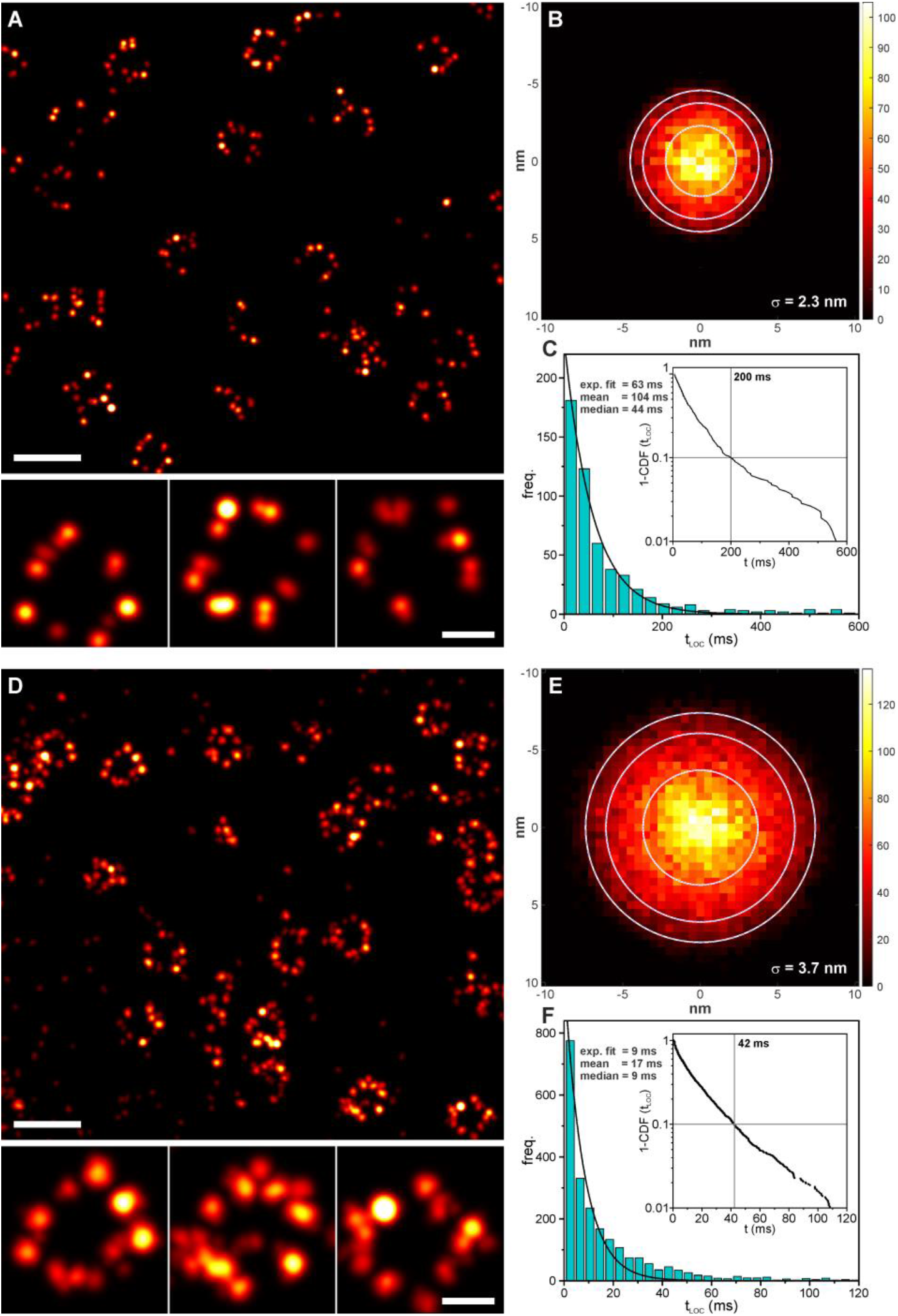
MINFLUX images of mEGFP-Nup107 cells labeled with nanobodies tagged with dye **4** (DOL = 2), with a 6 ms/iteration “slow” sequence (A), and with a 1.2 ms/iteration “fast” sequence (D). Selected individual NUPs are shown at the bottom of the panels. (B, E) 2D dispersion plots (a distance between the localization and the mean position of the cluster) were fitted with a 2D Gaussian, and mean sigma value is indicated, along with circles of 1x, 1.64x, and 2xσ. (C, F) Histograms for T_LOC_ (Δt = t_LAST_ – t_FIRST_) are also presented, with the experimental cumulative distribution function (the indicated value corresponds to 90% of the events).

## CONCLUSIONS

We introduced a series of red-emitting, stochastically blinking silicon rhodamine amides, derived from the widely used SiR fluorophore by the introduction of a thiophene- and benzothiophene-fused spirolactam fragment. The properties of the new fluorophores can be easily optimized by chemical modification of the pendant *N*-aromatic amide substituent. We performed a systematic characterization of the fluorophores at the single molecule level, mimicking the conditions of fluorescence nanoscopy experiments. We observed variations of the photophysical behavior of the dyes according to the type of chemical and biomolecular tag used, as well as a notable non-exponential behavior of the on-times affecting duty cycle distributions. With all six tested fluorophores, imaging using coordinate-stochastic super-resolution methods based on widefield (camera-based) single molecule localization was possible in fixed cells, and HaloTag ligands were found suitable for live-cell labeling. Remarkably, the new blinking fluorophores afforded considerably shorter on-times compared to those provided by most blinking fluorophores reported in the literature. With the best fluorophore versions, MINFLUX microscopy with down to 2.3 nm localization precision was achieved using HaloTag ligand and anti-GFP nanobodies labeled with a hydrophilized thiol-reactive maleimide derivative. With only a small trade-off in precision (3.7 nm) with respect to previous work, namely by a factor of 4, we accelerated the localization on a commercial MINFLUX instrument up to 30-times. We believe the strategy presented in this work will prompt the development of spontaneously blinking fluorophores optimized for MINFLUX imaging and stimulate further adaptation and optimization of MINFLUX imaging procedures to the needs and demands of future fluorophore families and their specialized nanoscopy applications.

## MATERIALS AND METHODS

### General Methods

Absorption and emission spectra as a function of pH were measured with a CLARIOstar Plus microplate reader (BMG LABTECH GmbH, Germany) in 96-well non-binding polystyrene F-bottom, μClear microplates (200 μL/well; Greiner Bio-One GmbH, Ref. 655906) at 25 °C in air-saturated solutions in triplicates. For determination of p*K*_a1_ and p*K*_a2_ values, plots of the absorption at λMAX (in the visible range) vs. p*H* were fitted to the following equation:

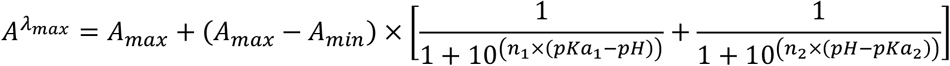

Fluorescence quantum yields were obtained with a Quantaurus-QY absolute PL quantum yield spectrometer (model C11347-11, Hamamatsu) according to the manufacturer’s instructions.

### Protein labeling

Polyclonal antibodies (goat anti-rabbit IgG (H+L), table 1) were labeled using a standard procedure. The pH of 2 mg/ml solutions was adjusted to ≈8.0 with a 1 M sodium bicarbonate solution and stirred for 1h at r.t. with 10 equiv of the NHS ester for STORM imaging, and 1 equiv for single molecule experiments. The mixture was purified using size-exclusion chromatography (PD MidiTrap G-25, GE Healthcare Bio-Sciences, gravity protocol) and stored at 4°C. The degree of labeling (DOL) obtained was calculated from spectrophotometric measurements as ≈2-3, for the concentrated mixtures, and estimated as ≈0.2-0.3 for the diluted mixtures from the reactive ratios. Anti-GFP nanobodies were labeled with compound **4-Maleimide** using a previously described protocol,^65^ and the DOL corroborated by mass spectrometry (Figure S10A). Binding of the HaloTag ligand derived from compound **4** *in vitro* was corroborated by mass spectrometry (Figure S10B). A mixture of 1 µM of the dye was mixed in PBS with 1.05 equiv. of the protein, incubated for 2 h at r.t., and submitted for mass analysis.

### Single Molecule Samples

Samples were prepared by a modified procedure (Figure S3), based on reported immobilization methods.^66, 67^ Flow chambers were constructed by attaching an oxygen plasma cleaned coverslip to an objective slide with double-sided adhesive tape. The chamber was incubated with PLL-PEG-biotin (0.2 mg/ml) and Tween-20 (1% v/v) in ddH_2_O for 15 min, rinsed with PBS, incubated with Streptavidin (10µg/ml) in PBS for 5 min, rinsed with PBS, incubated with biotinylated polyclonal antibody (≈10 µg/ml) from rabbit for 5 min, rinsed with PBS, incubated with dye-labeled polyclonal anti-rabbit antibody (≈10 µg/ml) for 5 min, rinsed with PBS, and filled with PBS for measurements. For HaloTag assays, the same chambers were incubated with PLL-PEG-NTA (0.2 mg/ml) and Tween-20 (1% v/v) in ddH2O for 15 min, rinsed with PBS, incubated with NiCl_2_ (2µg/ml) for 5 min, rinsed with PBS, incubated with his-tagged HaloTag7 (≈0.5 µM), rinsed with PBS, incubated with HaloTag ligand of the corresponding dye (10 nM), rinsed with PBS, and filled with PBS for measurements. The used reagents are summarized in Table S4.

### Cell Culture and Fluorescence Labeling

COS-7, HK-2xZFN-mEGFP-Nup107 and U2OS-Vim-Halo cells were cultured in Dulbecco’s modified Eagle medium (DMEM, 4.5 g/l glucose) in a CO_2_ incubator humidified at 5% at 37°C. The medium contains GlutaMAX and sodium pyruvate (ThermoFisher 31966), and supplemented with 10% (v/v) fetal bovine serum (FBS, ThermoFisher 10500064) and 1% penicillin-streptomycin (Gibco 15140122). U2OS-Nup96-Halo cells were cultured in McCoy’s 5A medium (Gibco 26600023) in a CO_2_ incubator at 37°C humidified to 5%. The medium contains L-glutamine and sodium pyruvate and is supplemented with 10% (v/v) fetal bovine serum (FBS, ThermoFisher 10500064) and 1% penicillin-streptomycin (Gibco 15140122). The cells were plated on glass coverslips for 24-48 h prior to fixing or labeling.

Halo cell lines were labeled live with the HaloTag ligands (250 nM) for 24 h in the corresponding medium, and washed twice with medium for at least 1 h. For live-cell imaging, samples were mounted in FluoBrite (Invitrogen) supplemented with 10% FBS (ThermoFisher) and 2% GlutaMAX (Gibco). For fixed-cell imaging (U2OS-Nup96-Halo), samples were washed with PBS, and fixed with PFA (3%) at room temperature for 20 min, quenched with 0.1 M NH_4_Cl and 0.1 M glycine in PBS (QB) for 10 min, permeabilized with Triton X-100 0.1% in PBS for 5 min and washed twice with PBS (5 min). For counter-labeling, they were stained with WGA AF488 in PBS and washed twice with PBS. mEGFP cells were fixed, permeabilized and blocked according to a reported protocol,^53, 68^ and labeled with the self-blinking nanobody conjugate for 60 min at r.t. COS-7 cells were washed with PBS, fixed for 5 min with MeOH cooled to −20°C, rinsed twice with PBS and blocked with BS for 45-60 min. Cells were incubated with a primary rabbit antibody against α-tubulin for 60 minutes at r.t. (or overnight at 4°C) in BS, washed twice with BS for 5 min, and then incubated with the anti-rabbit conjugate of the dye of interest in BS for 60 min at r.t. Afterwards samples were washed with BS, then twice with PBS.

All fixed samples were mounted on a concave coverslide filled with PBS, and sealed with a two-component silicon resin (Picodent Twinsil, Picodent Dental-Produktions-und Vertriebs-GmbH). Details on the used antibodies, nanobodies and proteins are summarized on Table S5.

### Single Molecule Characterization Setup

Characterization was performed in a custom-built microscope setup that can be switched between wide-field and confocal mode. A scheme with a more detailed description can be found in Figure S14. The microscope is equipped with a laser for excitation at 642 nm (MPB Communications, *P*_*max*_ = 1000 mW) and a laser for optional activation at 405 nm (Cobolt MLD-06, Hübner). The sample is placed on a custom-built stage based on a 3 degrees of freedom piezo positioning system (XYZ-SLC1740, SmarAct), which is built on a commercial Leica DMi8 (Leica Microsystems, Wetzlar, Germany) microscope. Illumination of the samples in both modes is done through a Leica HCX PL APO NA 1.46 Oil corrected objective lens. For wide-field detection an EMCCD camera is used (iXon DVU897, Andor Oxford Instruments, Abingdon UK), which detects light according to the filters built in the microscope body (660 nm, SR HC 660). For confocal detection, emission light is focused onto an avalanche photo diode (Excelitas SPCM-AQRH-13, Excelitas Technologies Corp. Waltham, USA), detecting light with a wavelength between 660 nm and 800 nm (Semrock 731/137 Brightline HC). The setup is controlled with a custom-written LabView software, which communicates with single components via a field programmable gate array board (FPGA, PCIe-R7852r, National Instruments, Austin, USA). Operating at a base clock frequency of 100MHz, it allows real-time control of the data acquisition. Wide-field data is acquired with the aid of the software provided by the manufacturer (Andor Solis), which interacts with the LabView software.

### Single Molecule Characterization

Single molecule data was acquired with any of the two illumination techniques described above and analyzed with custom-written MATLAB (MathWorks, USA) scripts. Separation of the signal from the noise was done by setting first an approximated threshold well above the background noise. Then, the signal below this threshold was fitted with a Gaussian (wide-field measurements) or a Poissonian (confocal measurements) distribution function. A final threshold was calculated excluding 99,999% (Gaussian distribution) and 99,994% (Poissonian distribution) of the noise. Then, single molecule traces were binarized using that threshold and most relevant parameters were calculated from the binary curves (e.g. T_ON_, T_OFF_, DC, N_C_), except frequencies, total photons and photons/cycles, which were calculated from the product of the trace and its binarized trace. Figure S5 shows a scheme visualizing the used method. The number of cycles (N_C_) is obtained by counting the number of positive flanks in the binary trace. The total number of photons (N_P_) was calculated as the sum of counts of the product of the trace and its binarized trace. The mean value of photons per cycle (PH_CY_) was calculated from the ratio of the previous values (N_PH_ / N_C_). To evaluate average on- and off-times, the histogram of all calculated events was plotted (Figures S5-S7) and fitted to either a mono-exponential function (T_OFF_) or a bi-exponential function (T_ON_). The average photon rate can be obtained by dividing the photons/event (N_PH_) through the duration of the event (T_ON_).

### STORM Imaging

STORM images were acquired on the setup described for single molecule characterization, on the wide-field mode, but excitation was performed with HILO (“highly inclined and laminated optical sheet microscopy”,^69^). Typically, 5000 frames for live-cell images, and 30000 frames for fixed cells were recorded. Data analysis was performed using ThunderSTORM ImageJ plugin^70^, including data merging and drift correction. Data was further filtered for outliers based on the sigma parameter (localization Gaussian fitting) and the uncertainty, and rendered by a custom -written MATLAB script. The images were rendered as Gaussians with a fixed size corresponding to the mean uncertainty of the localizations.

### MINFLUX Imaging

MINFLUX images were acquired on a commercial Abberior 3D MINFLUX microscope, equipped with a 560 and a 640 nm (cw) excitation laser line. The system also contains a 488 nm laser line only for confocal imaging, and a 405 nm laser for activation. In our measurements, only the excitation line at 640 nm was used. Detection was performed in two channels with the ranges 650 – 685 nm and 685 – 720 nm, respectively; signal from both channels was added up. The system is also equipped with a real-time position stabilization system, based on an infrared laser (975 nm) and a wide-field detection system of the light scattered at the sample interface. Image acquisition was done with different excitation powers starting from 240µW – 540µW (in front of the scanner) at the MINFLUX localization step. Different distances (*L*) for the last MINFLUX step have been taken depending of the aim of the measurement. Two imaging sequences were used for imaging, the standard (slow) and the optimized (fast) one. Table S3 resumes the main steps and imaging parameters.

Image and data post-processing was performed by custom-built MATLAB routines, described previously.^35^ Briefly, the data was first filtering with a density-based clustering algorithm (DBSCAN, epsilon = 4-8 nm, minPts = 3). Then, events with low emission frequency (<35kHz-55kHz depending on the sequence), with outside-inside ratio (CFR) of the photon count outside the range -0.5 – 0.8, and with less than 2 localizations were filtered out. Negative CFR values are assumed to come from center frequencies that are slightly below the average background level. A second filter was applied based on the spread of the localizations around their mean value. The distribution was first fitted with a 2D-Gaussian function, and localizations lying outside a radius corresponding to 90% (1.645 × *σ*) of the distribution were discarded. The images were then plotted via an amplitude-normalized Gaussian rendering (pixel-size = 1 nm) with fixed sigma (σX= σY) as the average value from the distribution of all remaining (filtered) molecules. For visualization purposes (the number of localizations varies with on-time and number of cycles) the color maps used in the images are non-linear with a gamma correction of *A* = 1 and *γ* = 0.5. We further extracted the number of photons per localization, the total number of photons from a trace (N_PH_, including the pre-localization and all localizations), and the localization time (T_LOC_) calculated from the difference in time-stamp of its last and first localizations. The mean, the median and the expectation value (τ) from a mono-exponential fit of the distributions (histogram) of each variable were calculated.

## AUTHOR CONTRIBUTIONS

A.N.B. and M.L.B. were responsible for the project’s conception. M.L.B. and M.R. wrote the manuscript with input from A.N.B. and S.W.H. A.N.B. performed the chemical synthesis; M.R., L.S., M.L.B. and A.N.B. performed dye characterization; M.R. and M.L.B. performed labeling, microscopy, and data analysis; S.W.H. directed and supervised the investigations; all the authors discussed the results and commented on the manuscript.

## ACKNOWLEDGMENT

A.N.B. and M.L.B. acknowledge funding from the German Federal Ministry of Education and Research (BMBF Project 13N14122 “3D Nano Life Cell”, to S.W.H.). We thank Prof. S. Jakobs (MPI for Multidisciplinary Sciences, Göttingen) for providing the U2OS-Vim-Halo cells and the European Molecular Biology Laboratory (Heidelberg) for providing the U2OS-NUP96-Halo and HK-2xZFN-mEGFP-Nup107 cells. We also thank the following co-workers at the MPI for Medical Research (Heidelberg): Dr. S. Fabritz and the staff of Mass Spectrometry Core Facility for recording mass spectra of proteins and small molecules, M. Steigleder (Electronics Workshop) for building custom electronics for data acquisition, Dr. M. Tarnawski and the staff of the Protein Expression and Characterization Facility for providing the HaloTag7 protein, Dr. E. D’Este (Optical Microscopy Facility) for the access to MINFLUX microscope and its maintenance, J. Hubrich and A. Fischer for cell culture and tips on labeling, Dr. J. Engelhardt for assisting with the construction of the single molecule characterization setup, Dr. J. Matthias for input on cell labeling, and O. Wolff for fruitful discussions on statistics and coding. We acknowledge Dr. M. Weber (MPI for Multidisciplinary Sciences) and Dr. L. J. Patalag (Technische Universität Braunschweig) for the preliminary evaluation of spontaneously blinking fluorophores.

