## Supplemental figures and text. for "Spontaneously blinking fluorophores for accelerated MINFLUX nanoscopy"

### "Spontaneously blinking fluorophores optimized for fast localization with MINFLUX nanoscopy"

#### Supplementary Figures:

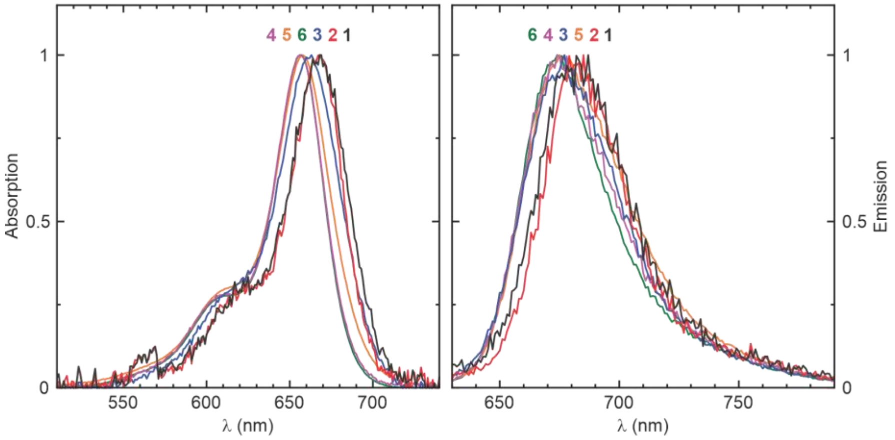

**Figure S1.** Normalized absorption and emission spectra of compounds **1-6** in a buffered solution at pH = 7, corresponding to the open xanthylium form (see Figure 1).

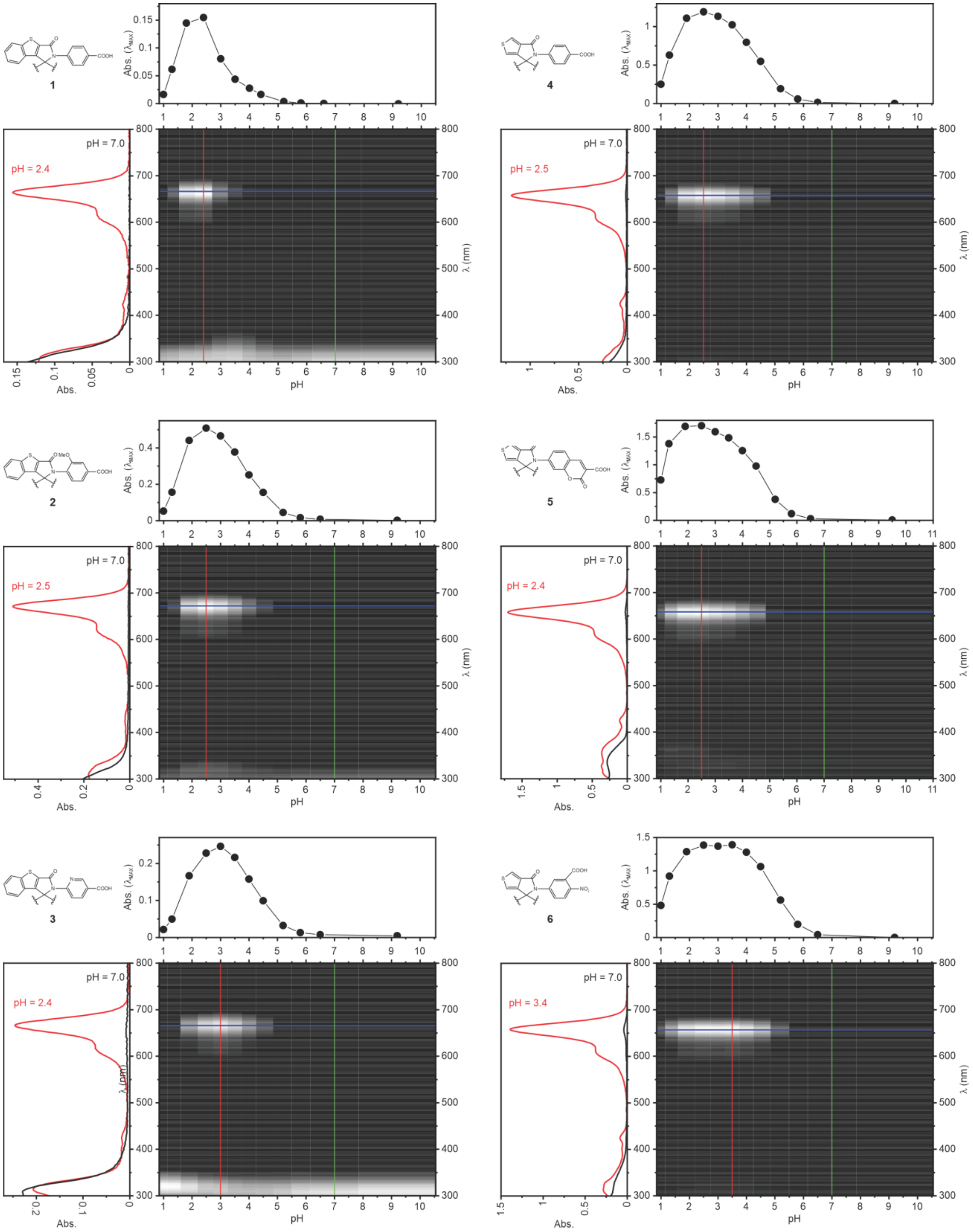

**Figure S2.** 2D absorption/pH maps (center) for compounds **1-6** (10 µM). The absorption changes at the absorption maximum (top) and the spectra at pH = 7 and at the pH where maximum absorption is reached (left) are also shown.

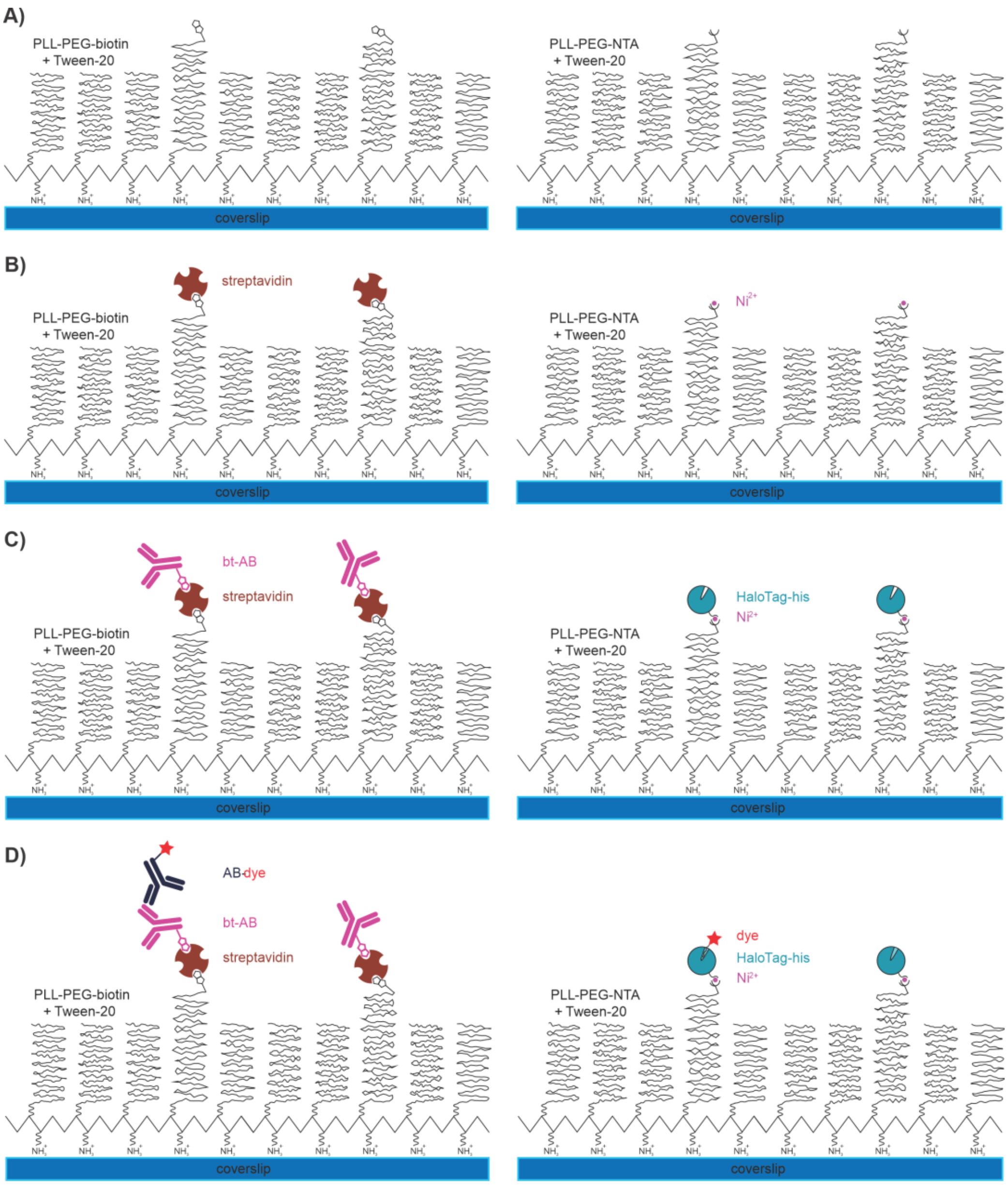

**Figure S3.** Schematic representation of the protocol uses for the preparation of samples for single molecule characterization experiments. Samples for the in vitro antibody studies were prepared by (A) coating a plasma-cleaned coverslip with PLL-PEG-biotin co-polymer with Tween-20 added to minimize unspecific binding of protein [1]. (B) Streptavidin was used to immobilize (C) primary biotinylated antibodies (bt-AB). (D) Incubation of a secondary antibody labeled with the fluorophore gave sparsely labeled single molecule samples. Samples for the in vitro HaloTag studies were prepared by (A) coating a plasma-cleaned coverslip with PLL-PEG-NTA co-polymer with Tween-20 added to minimize unspecific binding of protein [2]. (B) After NTA groups were loaded with $Ni^{2+}$, (C) HaloTag7 proteins (his-HT7) were immobilized with their $his_6$-tags. (D) Incubation of fluorescent HaloTag ligand gave sparsely labeled single molecule samples. Specific binding in both samples was confirmed by a drastically reduced immobilization in negative controls that missed bt-AB or his-HT7, respectively.

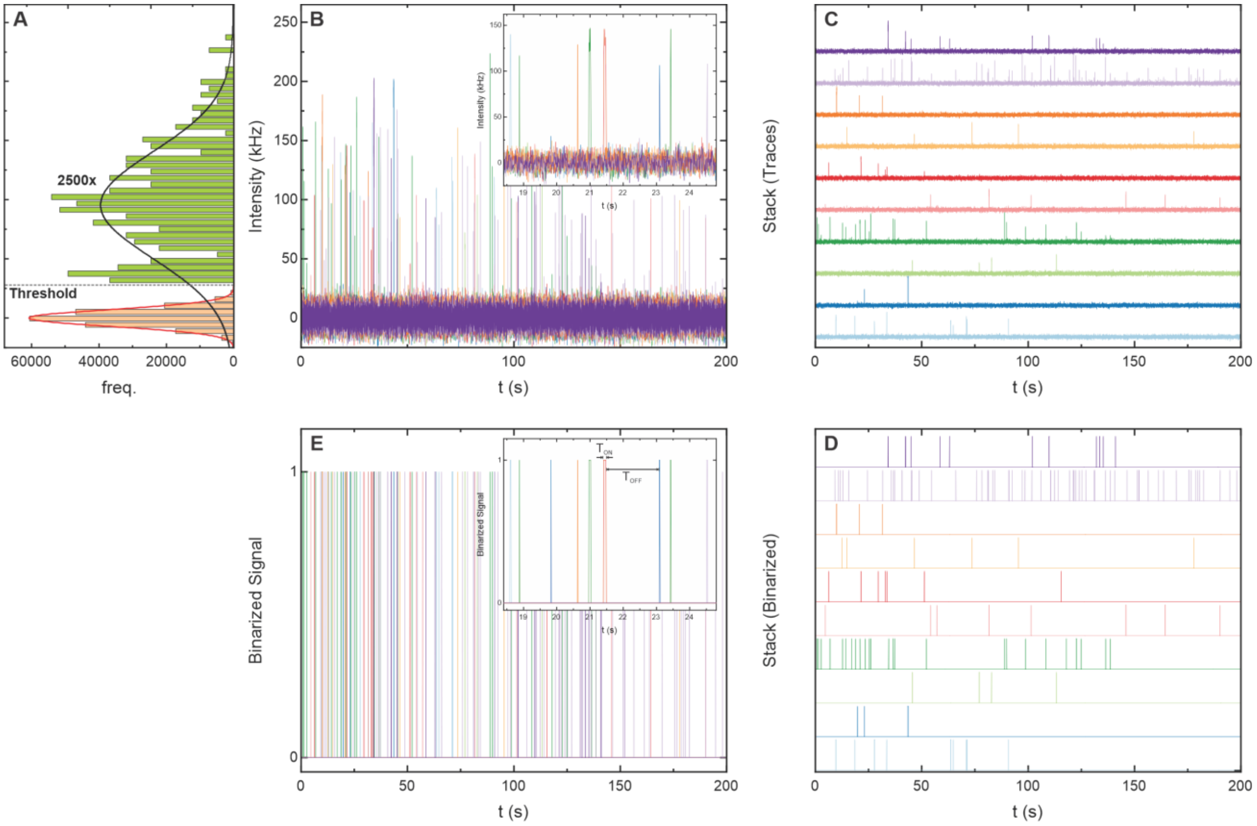

**Figure S4.** Example of data analysis for the extraction of $T_{ON}$ and $T_{OFF}$ from single molecules traces. On the upper side, ten single molecule traces are shown overlaid (B) and stacked (C). On the left side (A) the intensity histograms are display with the selected threshold; resulting noise and single molecule events are shown at a different scale and color. The threshold was calculated to include in the noise 99,999% (99,994 for poissonian) according to the fitted distribution. The separation allows plotting binarized traces (E-D), which were used to calculate $T_{ON}$, $T_{OFF}$, DC, $N_C$. Total photons and the emission rate were calculated from the original traces.

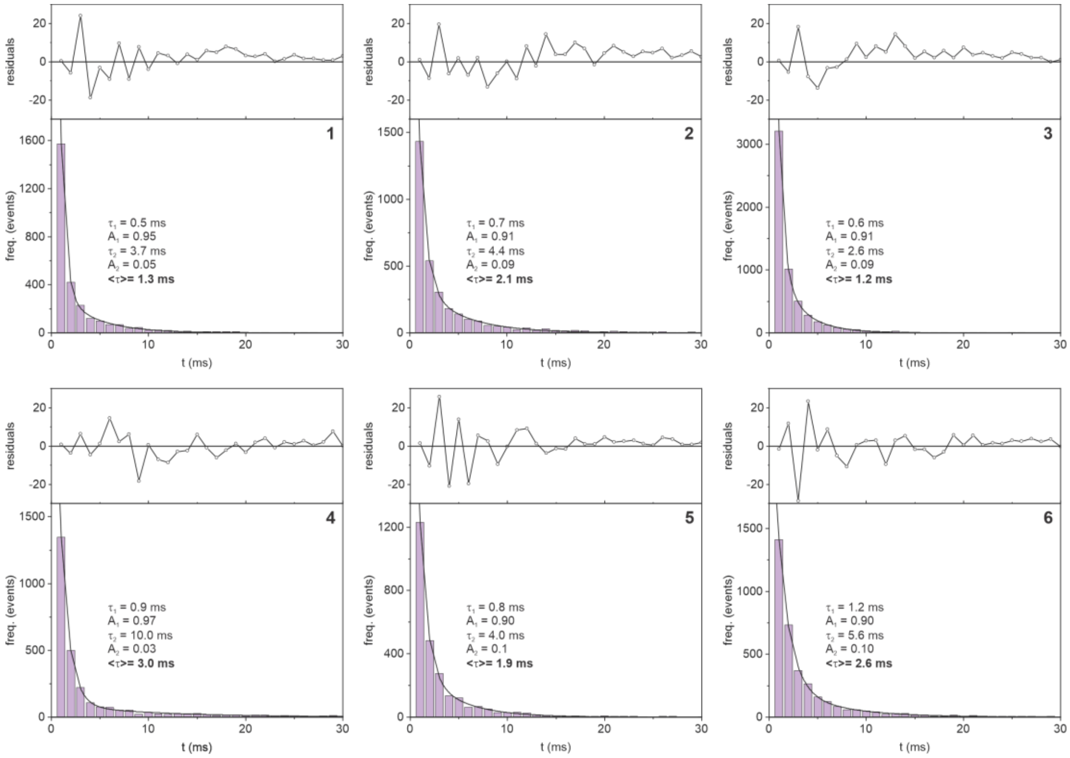

**Figure S5.** Histograms for the on-times ($T_{ON}$), with a bi-exponential fit (lines) and residuals (top plots), for compounds **1-6**. Fitted parameters and the calculated average on times are indicated on each plot.

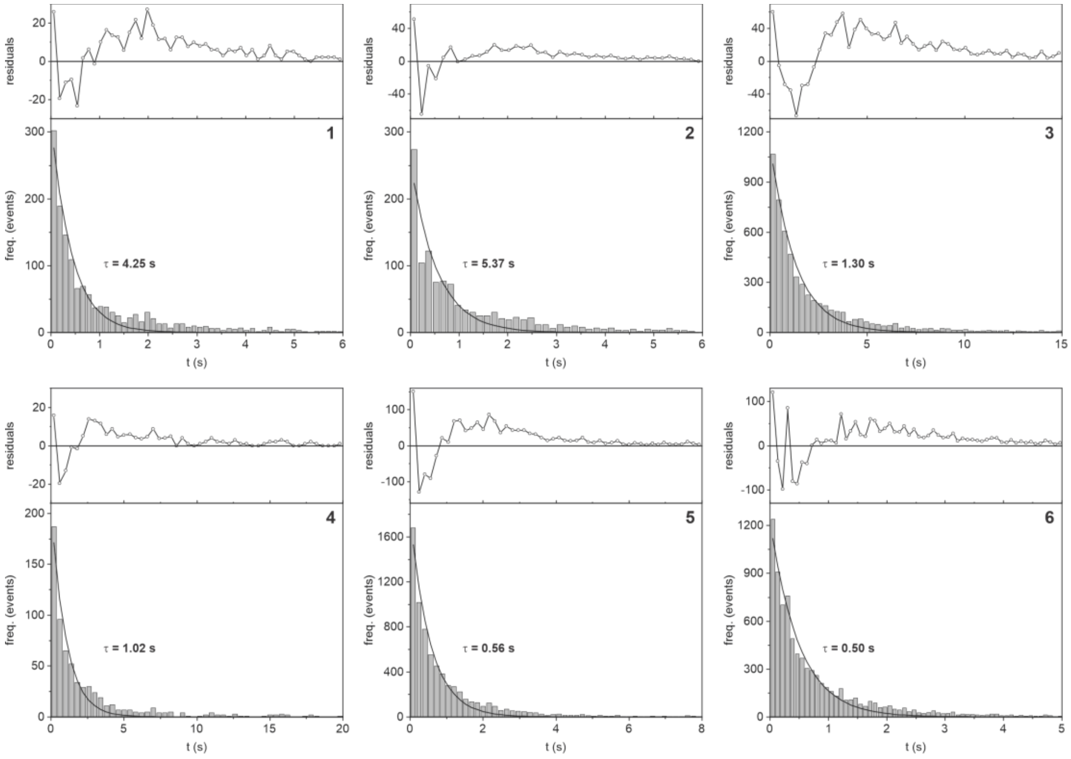

**Figure S6.** Histograms for the off-times ($T_{OFF}$), with a mono-exponential fit (lines) and residuals (top plots), for compounds **1-6**. The fitted times are indicated on each plot.

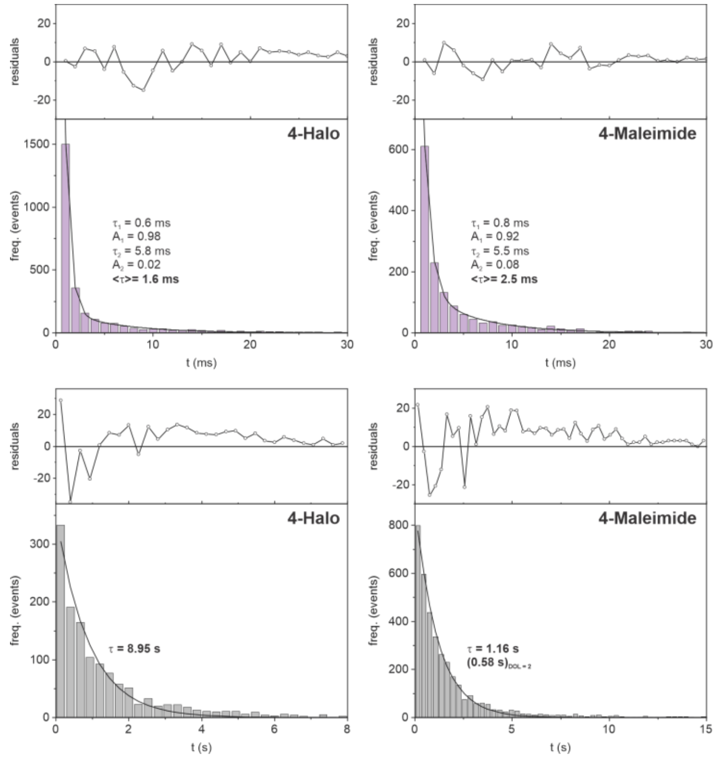

**Figure S7.** Histograms for the on- ($T_{ON}$) and off-times ($T_{OFF}$), with the corresponding fits (lines) and residuals (top plots), for compound **4-Halo** bound to HaloTag7 protein, and a nanobody labeled with compound **4-Maleimide**. Fit parameters and the calculated average on-times are indicated on each plot. Note that the off-time of the nanobody adduct is halved from the fitted value, to consider the DOL = 2 (each protein contains two blinkers). Calculated duty cycles are $1.82\times10^{-4}$ (HT) and $2.15\times10^{-3}$ (NB).

**Table S1.** Photophysical properties for compound **4** on antibodies (NHS), **4-Halo** bound to HaloTag7 protein, and **4-Maleimide** adduct with nanobodies. The duty cycle is calculated as $\mathrm{DC} = \frac{\mathrm{T_{on}}}{\mathrm{T_{on}}+\mathrm{T_{off}}}$ with $\mathbf{T_{on}}$ acquired from confocal illumination and $\mathbf{T_{off}}$ acquired from wide-field illumination. The number of cycles $\mathbf{N_{cy}}$ are the mean on events calculated per molecule. As the dye might not be bleached by the end of the measurement (100s exposure) this has to be seen as lower limit. **%Bl** was obtained by calculating a theoretical $\mathbf{N_{cy,theo}}$ and comparing it with the measured $\mathbf{N_{cy}}$: $\%\mathrm{Bl} = \frac{\mathrm{N_{cy}}}{\mathrm{N_{cy,theo}}} = \frac{\mathrm{N_{cy}}}{\mathrm{T_{off}} \cdot \mathrm{N_{frames}}}$. The photons/cycle were estimated as $\mathrm{Ph_{cy}} = \frac{\mathrm{Mean(photon)}}{\mathrm{mean(N_{Cy})}}$, and the rate $\mathrm{Rate} = \frac{\mathrm{Ph_{cy}}}{\mathrm{T_{on}}}$.

| Comp | DC X1000 | $T_{ON}$ / ms | $T_{OFF}$ / s | $N_{CY}$ | %Bl | $PH_{CY}$ | Rate (kHz) |
|---|---|---|---|---|---|---|---|
| **Antibody** | 2,89 | 3,0 | 1,02 | 14 | 86 | 354 | 118 |
| **Halotag** | 0.18 | 1.6 | 8.95 | 6.1 | 45 | 175 | 109 |
| **Nanobody** | 2.15 | 2.5 | 1.16 | 6.1 | 86 | 184 | 73 |

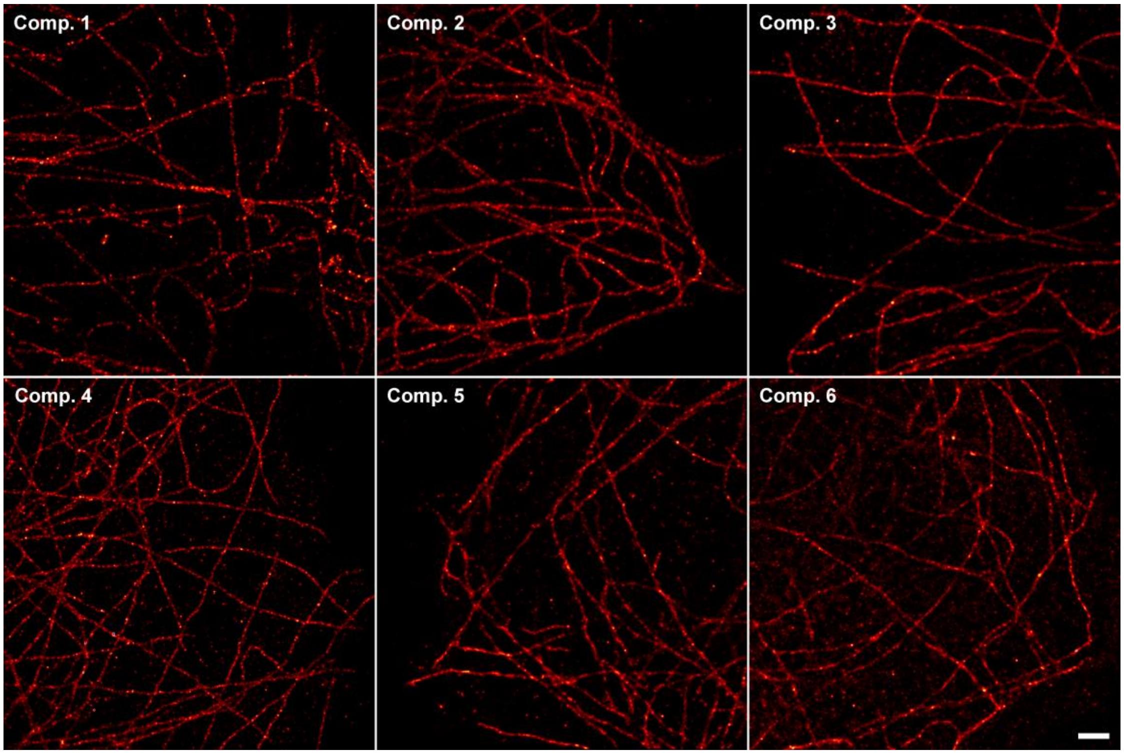

**Figure S8.** STORM imaging on fixed cells, stained with a primary antibody (anti-tubulin) and secondary antibodies labelled with compounds **1-6** (high DOL). Scale bar (1 µm)

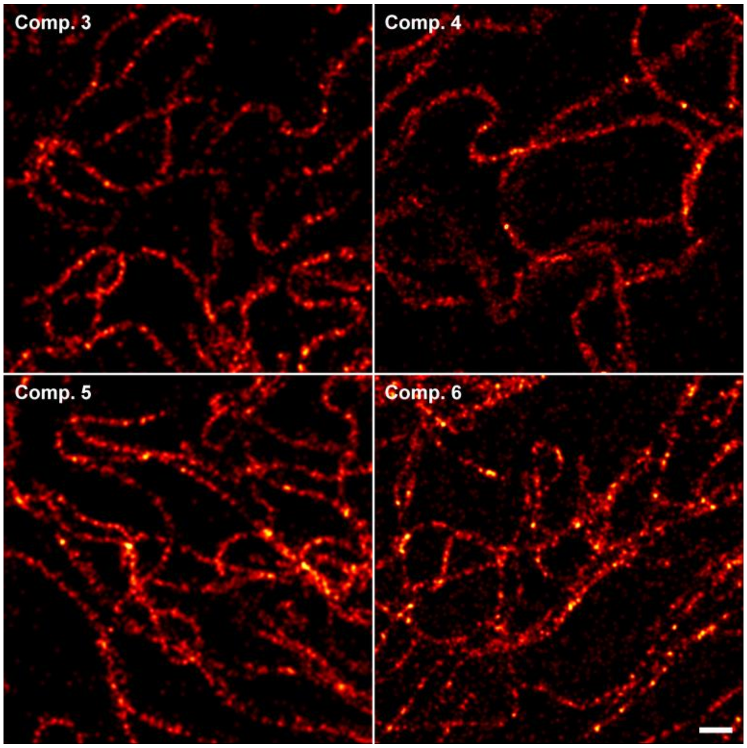

**Figure S9.** STORM imaging on live-cells, labeled with chloroalkane ligands (vimentin) of compounds **3-6**. Scale bar (500 nm)

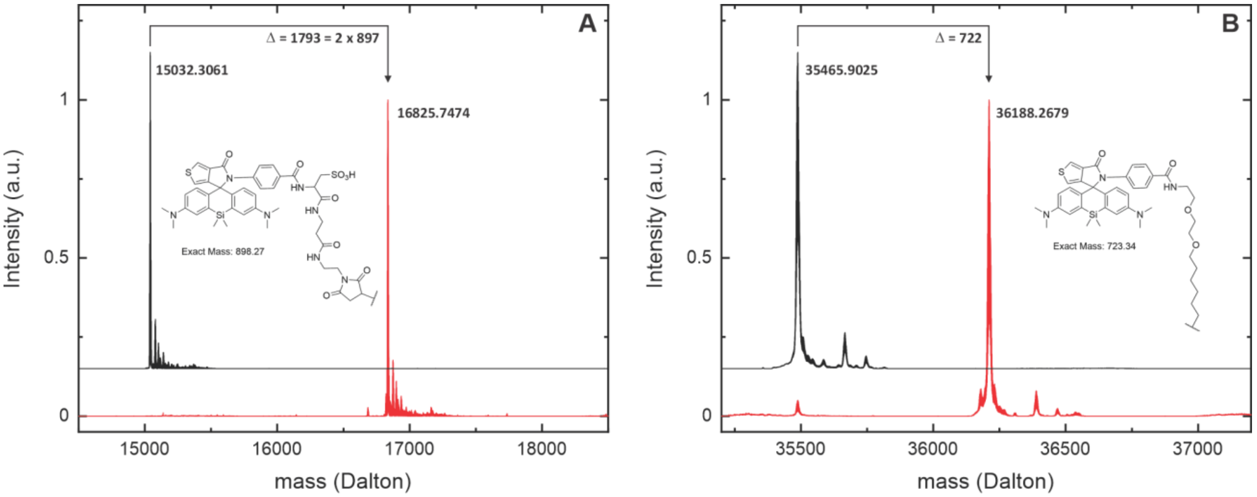

**Figure S10.** (A) Mass spectra of the unlabeled nanobody (black line) and labelled with **4-Maleimide** (red line), after purification red curve); (B) Mass spectra of free HaloTag7 protein (black) and HaloTag7 labeled with **4-Halo** (5-8% mol excess of protein, > 1 h at rt), without purification.

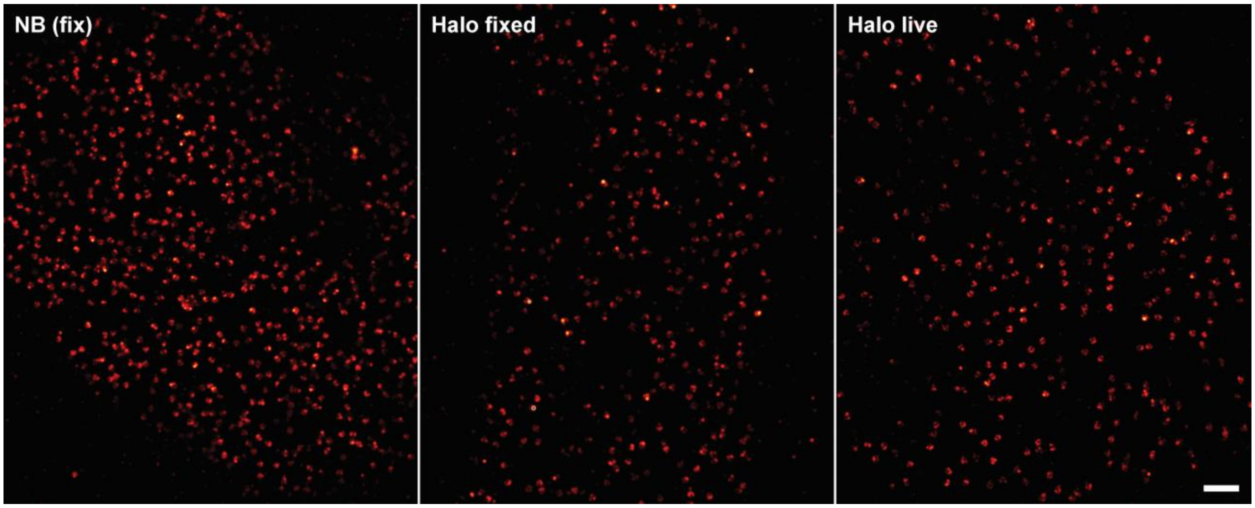

**Figure S11.** STORM imaging on fixed and live-cells with compound **4** (**4-Maleimide** or **4-Halo**), labeled via nanobodies (fixed), HaloTag (live labelling, then fixed before imaging), and HaloTag (live imaging). Fixed samples were mounted in PBS and live samples were mounted in supplemented FluoBrite cell medium (Invitrogen). Scale bar (1 µm)

**Table S2.** Main parameters used in the MINFLUX imaging sequences. MINFLUX *L* parameter is the characteristic distance of the target coordinate excitation pattern (distance between center and outside expositions); Center frequency ratio (*cfr*) is the photon ratio between center and outside exposures; the dwell time (*dt*) is the minimal time of one step in which the photon threshold (*thr*) has to be surpassed; the offset background (BG) is a frequency threshold an event must surpass to be accepted; the power factor (*PF*) is a multiplier of the base excitation intensity set for the first step.

| | slow sequence | | | | | | fast sequence | | | | | |
|---|---|---|---|---|---|---|---|---|---|---|---|---|
| | *L* (nm) | *Thr* (phot) | *cfr* | *Dt* (ms) | offset BG (kHz) | *PF* | *L* (nm) | *Thr* (phot) | *cfr* | *Dt* (ms) | offset BG (kHz) | *PF* |
| step 1 (gauss) | 288 | 30 | 2.0 | 1 | 10000 | 1 | 288 | 20 | 1 | 0.4 | 15000 | 2 |
| step 2 (donut) | 288 | 30 | 0.5 | 1 | 8000 | 1 | 151 | 30 | 0.8 | 0.3 | 10000 | 2 |
| step 3 (donut) | 151 | 30 | 2 | 1 | 8000 | 2 | 76 | 30 | 0.8 | 0.5 | 30000 | 4 |
| step 4 (donut) | 101 | 30 | 0.8 | 1 | 8000 | 4 | | | | | | |
| step 5 (donut) | 76 | 30 | 0.8 | 1 | 8000 | 4 | | | | | | |
| step 6 (donut) | 40 | 30 | 0.8 | 1 | 8000 | 6 | | | | | | |

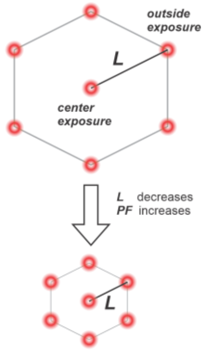

**Table S3.** Localization parameters obtained from MINFLUX images shown in Figure 4 and 5.

| | Binning (photons) | σ / nm | $N_{LOC}$ (median/exp) | $N_{PH}$ (median) | $T_{LOC}$ / ms (median/exp) | $Phot_{Preloc}$ (median) | $Phot_{Loc}$ (median) |
|---|---|---|---|---|---|---|---|
| NB (4A) | --- | 2.6 | 7/10 | 1300 | 52/70 | 1000 | 51 |
| HT (4C) | --- | 2.6 | 12/12 | 2600 | 103/136 | 1640 | 83 |
| Slow (5A) | --- | 2.3 | 7/9.4 | 2200 | 44/63.4 | 1800 | 71 |
| Fast (5B) | --- | 3.7 | 6/7.5 | 450 | 9/8.6 | 100 | 55 |
| Schmidt et al. | 350 | [2.2][a] / 1.35[b] | 5/3.9 | 2230 | 155/x | --- | 415 |
| (Figure 3d) [3] | 2100 | [0.9][a] / 0.8[b] | 3/--- | 6600 | 323/x | --- | 2173 |

[a] Reported in paper; [b] after applying the filtering method used in this work (*MATERIALS AND METHODS* section)

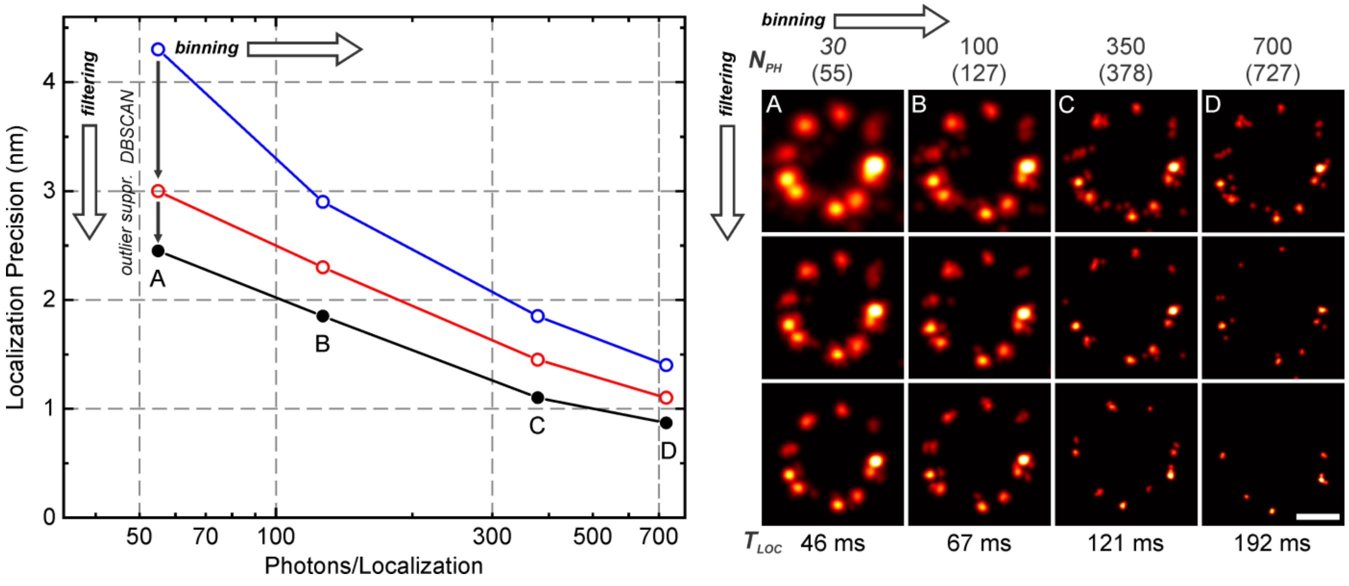

**Figure S12.** Localization accuracy as a function of binning, calculated for the image on Figure 4a with the raw data (blue symbols), and after applying a DBSCAN filter (red symbols) and the outlier suppression filter (black symbols). The data was binned by combining successive localizations until a total amount of photons ($N_{PH}$ = 100, 350, 700), on the last localization step, were reached. The starting value of 30 photons is set by the acquisition routine, and the values in brackets are the mean value calculated after binning. An image of a single NUP is shown on the right for each case. The average localization time ($T_{LOC}$), calculated for the filtered images, is indicated for each case. Moving from the top-left to lower-right results in a loss of used events due to a cutoff in photons (left to right), and to the uncertainty (top to bottom). In addition, molecules with short on-times are eliminated in both directions, as they are less likely to meet the corresponding criteria. Scale bar (50 nm).

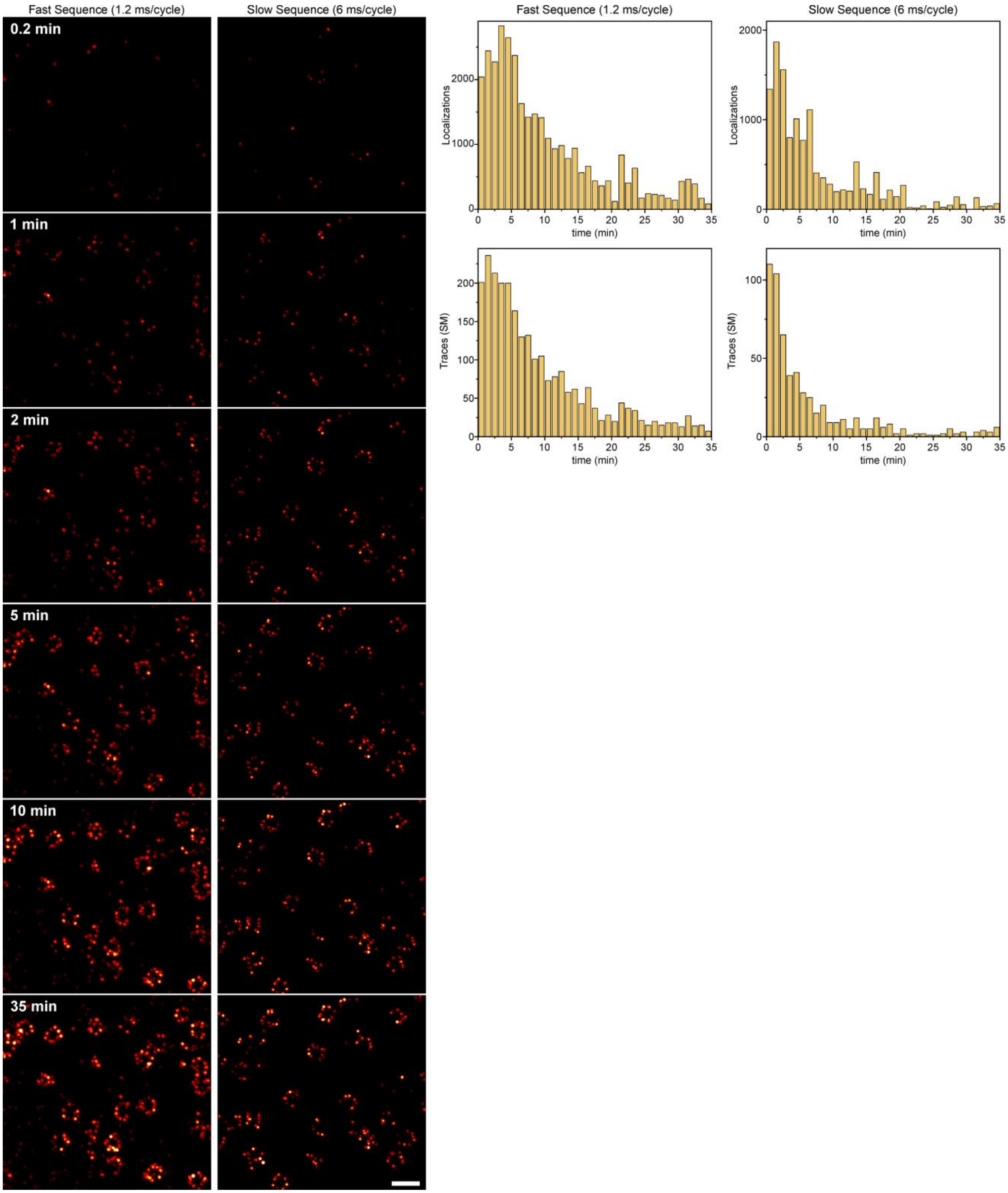

**Figure S13.** Dynamic image buildup from traces/localizations, for the images from Figure 5. Time distribution of localizations and traces (SMs) are also shown. Scale bar (200 nm).

**Table S4.** Proteins and additives used for single molecule characterization experiments.

| Reagent | Type | Supplier | Catalogue No. | Concentration |
|---|---|---|---|---|
| PLL-PEG-biotin (PLL(20)-g[3.5]-PEG(2)/PEG(3.4)- biotin(20%) | Polymer-Protein Layer | Suso AG Inc. | | 0.2mg/ml |
| PLL-PEG-biotin (PLL(20)-g[3.5]-PEG(3.4)-NTA, biotin(20%) | Polymer-Protein Layer | Suso AG Inc. | | 0.2mg/ml |
| Streptavidin | Protein | Merck/Sigma Aldrich | 189730 | 10µg/ml |
| $NiCl_2$ | Additive | Merck/Sigma Aldrich | 339350 | 2µg/ml |
| his-HaloTag7 | Protein | Protein Expresison Facility MPIMR | | 1:10000 |
| AffiniPure Goat Anti-Rabbit IgG (H+L) | Secondary antibody (goat, anti-rabbit) | Jackson ImmunoResearch Europe Ltd | 111-005-003 | 1:100 |
| chloroalkane dye adduct | Reactive dye | (prepared in this work) | | 10 nM |
| Biotin-SP (long spacer) AffiniPure Rabbit Anti-Mouse IgG (H+L) | Secondary antibody (rabbit, anti-mouse) | Jackson ImmunoResearch Europe Ltd | 315-065-045 | 1:100 |

**Table S5.** Antibodies and nanobodies used for labeling and imaging.

| Reagent | Type | Target | Host | Supplier | Catalogue No. | Dilution |
|---|---|---|---|---|---|---|
| Anti-α-Tubulin antibody | Primary Antibody (monoclonal) | α-tubulin | Rabbit | Abcam | ab18251 | 1:200 |
| Anti-Nup153 antibody | Primary Antibody (monoclonal) | Nup153 | Mouse | Abcam | ab24700 | 1:300 |
| FluoTag-X2 anti-GFP unconjugated clone 1H | Nanobody | GFP | Camelid | NanoTag Biotechnologies | N0302 | 1:4000 |
| Invitrogen Goat Anti-Mouse IgG (H+L) | Secondary antibody | Mouse | Goat | Thermo Fisher | A32723 | 1:1000 |
| chloroalkane dye adduct | Reactive dye | HaloTag | | (prepared in this work) | | 250nM |

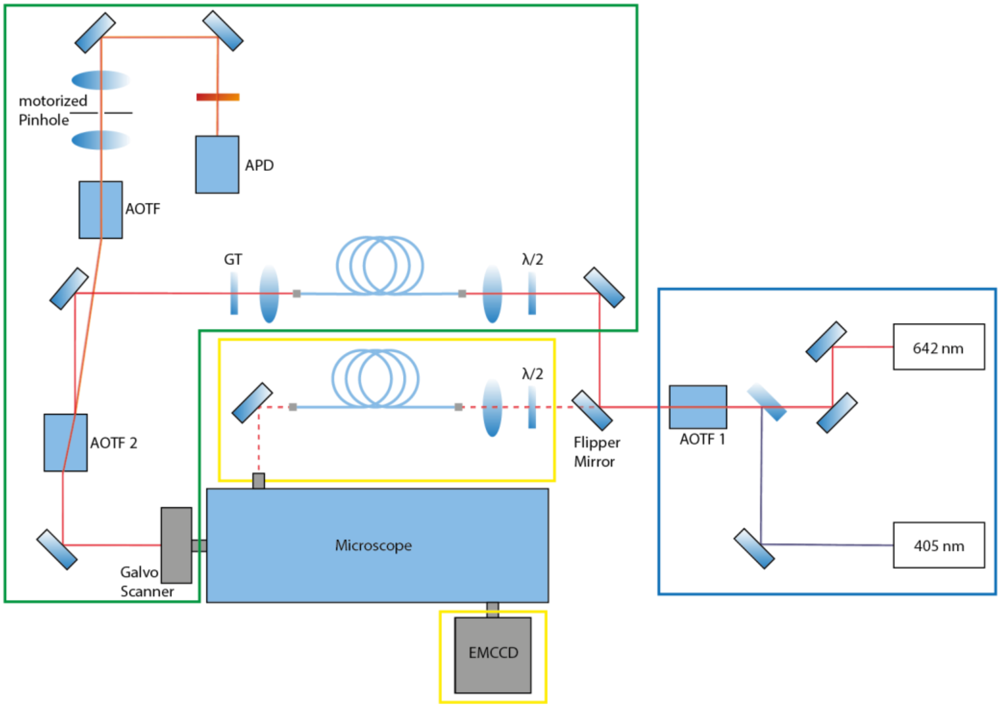

**Figure S14.** Schematic representation of the custom-built experimental setup used for the characterization of compounds (**1-6**) at the single molecule level, and to perform STORM/HILO imaging. Excitation light (642 nm) gets power adjusted by an acousto-optic-tunable-filter (AOTF 1, PCAOM VIS, Crystal Technology Inc.) in the excitation chamber (blue). With the aid of a flipper mirror (8892-K-M, Newport) light gets guided into the respective single mode polarization maintaining fibers (Thorlabs P1-405BPM-FC-5, Thorlabs Inc. Newton, USA) for either confocal (green) or wide-field (yellow) illumination. For confocal excitation, the polarization of the light exiting the fiber is clean up via a Glan-Thompson (GT) polarizer (Thorlabs GTH5M). After passing AOTF2, filtering everything but the excitation wavelength, it is coupled into an analog galvanometer scanner with four scanning units (mirrors: 6210H, servo driver: MicroMax™ 671, Cambridge Technology), which enables confocal scanning. The scanner guides the light into the body of a Leica DMi8 and is focused on the sample with the aid of a Leica HCX PL APO NA 1.46 Oil corrected objective lens. Fluorescence is collected via the same objective, de-scanned with the aid of the scanner and separated from the excitation light via AOTF 2. It passes a motorized pinhole refining the confocal volume imaged and filtered further by two dichroic filters (Semrock 731/137 Brightline HC). It's then focused on an avalanche photo diode. For wide-field illumination light exits the corresponding fiber and is coupled into the sideport of the microscope body via an adjustable dielectric mirror. The light gets focused into the pupil of the objective via the tube lens used in the side-port, which leads to a collimated beam exiting the objective. The dielectric mirror allows shifting of the focus spot in the pupil of the objective enabling HILO illumination. Separation of the excitation and emission light is done by a dichroic mirror inside the microscope body (660 nm, SR HC 660). Before getting focused onto the CCD chip of an EMCCD camera emission light is further filtered by a bandpass (665 – 732 nm, Chroma ET700/75). Realtime control of the setup is done by a self-written LabView software.

#### General experimental information and synthesis

**Thin layer chromatography:** Analytical TLC (normal phase) was performed on Merck Millipore ready-to-use aluminum sheets coated with silica gel 60 ($F_{254}$) (Cat. No. 1.05554.0001). Compounds were detected by exposing TLC plates to UV-light (254 or 366 nm) or by heating with vanillin stain (6 g vanillin and 1.5 mL conc. $H_2SO_4$ in 100 mL ethanol) or PMA stain (10 g phosphomolybdic acid hydrate in 100 mL ethanol).

**Preparative flash column chromatography:** Automated separations on normal phase were performed with an Isolera Spektra One system (Biotage AG, Sweden) using commercially available cartridges of suitable size (RediSep Rf series from Teledyne ISCO, Puriflash Silica HP 30µm series from Interchim) and solvent gradient as indicated for individual preparations.

**High-Performance Liquid Chromatography (HPLC):** Preparative high-performance liquid chromatography was performed on a Büchi Reveleris Prep system using Interchim 250×21.2 mm 5 µm Uptisphere Strategy PhC4 column and conditions as indicated for individual preparations. Method scouting was performed on a HPLC system (Shimadzu): 2x LC-20AD HPLC pumps with DGU-20A3R solvent degassing unit, CTO-20AC column oven equipped with a manual injector with a 20 µL sample loop, SPD-M20A diode array detector, RF-20A fluorescence detector and CBM-20A communication bus module; analytical column: Interchim 250×4.6 mm 5 µm PhC4, solvent flow rate 1.2 mL/min.

**Mass Spectrometry (MS):** Analytical liquid chromatography-mass spectrometry was performed on an LC-MS system (Shimadzu): 2x LC-20AD HPLC pumps with DGU-20A3R solvent degassing unit, SIL-20ACHT autosampler, CTO-20AC column oven, SPD-M30A diode array detector and CBM-20A communication bus module, integrated with CAMAG TLC-MS interface 2, FCV-20AH$_2$ diverter valve and LCMS-2020 spectrometer with electrospray ionization (ESI, 100 – 1500 m/z). Analytical column: ThermoScientific Hypersil Gold 50×2.1 mm 1.9µm, standard conditions: sample volume 1-2 µL, solvent flow rate 0.5 mL/min, column temperature 30 °C. General method: isocratic 95:5 A:B over 2 min, then gradient 95:5 to 0:100 A:B over 5 min, then isocratic 0:100 A:B over 2 min; solvent A – water + 0.1% (v/v) $HCO_2H$, solvent B – acetonitrile + 0.1% (v/v) $HCO_2H$.

High resolution mass spectra (HRMS) were obtained on a maXis II ETD (Bruker) with electrospray ionization (ESI) at the Mass Spectrometry Core facility of the Max-Planck Institute for Medical Research (Heidelberg, Germany).

**NMR spectra** were recorded at 25 °C with a Bruker Ascend 400 spectrometer at 400.15 MHz ($^1H$) and 100.62 MHz ($^{13}C$) and are reported in ppm. All $^1H$ spectra are referenced to tetramethylsilane (TMS; $\delta$ = 0 ppm) using the signals of added TMS (0.03% v/v) or the residual protons of $CHCl_3$ (7.26 ppm) in $CDCl_3$, $CHD_2CN$ (1.94 ppm) in $CD_3CN$,

$CHD_2COCD_3$ (2.05 ppm) for acetone-$d_6$, pyridine-$d_4$ (8.74 ppm, H-2, H-6) for pyridine-$d_5$. $^{13}C$ spectra are referenced to TMS ($\delta$ = 0 ppm) using the signals of added TMS (0.03% v/v) or the solvent: $CDCl_3$ (77.16 ppm), $\underline{C}D_3CN$ (1.32 ppm), $(\underline{C}D_3)_2CO$ (29.84 ppm), or pyridine-$d_5$ (150.35 ppm, C-2,6). Multiplicities of signals are described as follows: s = singlet, d = doublet, t = triplet, q = quartet, p = pentet, m = multiplet or overlap of non-equivalent resonances; br = broad signal. Coupling constants ($J$) are given in Hz. $CDCl_3$ solvent was freshly filtered before dissolving samples through a short plug of basic alumina.

#### Synthetic procedures for the preparation of fluorescent dyes

#### Compound S-1

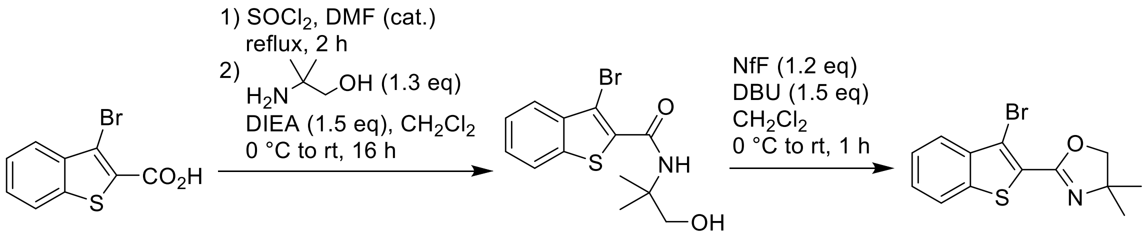

**S-1**

DMF (1 drop) was added to a suspension of 3-bromobenzothiophene-2-carboxylic acid (1 g, 3.89 mmol) in thionyl chloride (4 mL), and the mixture was refluxed for 2 h. The resulting yellowish solution was evaporated to dryness, chased twice with dry $CH_2Cl_2$ (5 mL), and the residue was dissolved in dry $CH_2Cl_2$ (5 mL). This solution was added dropwise to a stirred mixture of 2-amino-2-methyl-1-propanol (450 mg, 5.06 mmol, 1.3 equiv) and *N*,*N*-ethyldiisopropylamine (DIEA; 1.02 mL, 5.84 mmol. 1.5 equiv) in dry $CH_2Cl_2$ (10 mL), cooled in ice-water bath. The reaction mixture was allowed to warm up to rt and left stirring overnight (16 h). The amide product was extracted with $CH_2Cl_2$ (3 × 40 mL) for sat. aq. $NaHCO_3$ (50 mL), the combined extracts were washed with brine, dried over $Na_2SO_4$, filtered and evaporated to dryness.

The resulting crude amide was dissolved in $CH_2Cl_2$ (30 mL), 1,8-diazabicyclo[5.4.0]undec-7-ene (DBU; 0.87 mL, 5.84 mmol, 1.5 equiv) was added, the solution was cooled in ice-water bath followed by addition of perfluoro-1-butanesulfonyl fluoride (NfF; 0.84 mL, 4.67 mmol, 1.2 equiv). The reaction mixture was warmed up to rt and stirred for 1 h, then poured into sat. aq. $NaHCO_3$ (50 mL), extracted with $CH_2Cl_2$ (3 × 30 mL), the combined extracts were washed with brine and dried over $Na_2SO_4$. The product was isolated by flash column chromatography (40 g Teledyne ISCO RediSep Rf cartridge, gradient 5% to 30% EtOAc/hexane) to yield 1.12 g (93%) of **S-1** as viscous oil, which solidified in freezer overnight.

$^1H$ NMR (400 MHz, $CDCl_3$): δ 7.94 – 7.88 (m, 1H), 7.83 – 7.76 (m, 1H), 7.50 – 7.43 (m, 2H), 4.19 (s, 2H), 1.43 (s, 7H).

$^{13}C$ NMR (101 MHz, $CDCl_3$): δ 157.4, 138.9, 138.7, 127.3, 126.2, 125.6, 124.8, 122.5, 111.0, 79.8, 68.1, 28.4.

HRMS ($C_{13}H_{12}BrNOS$): *m/z* (positive mode) = 309.9892 (found $[M+H]^+$), 309.9896 (calc.).

#### Compound S-3

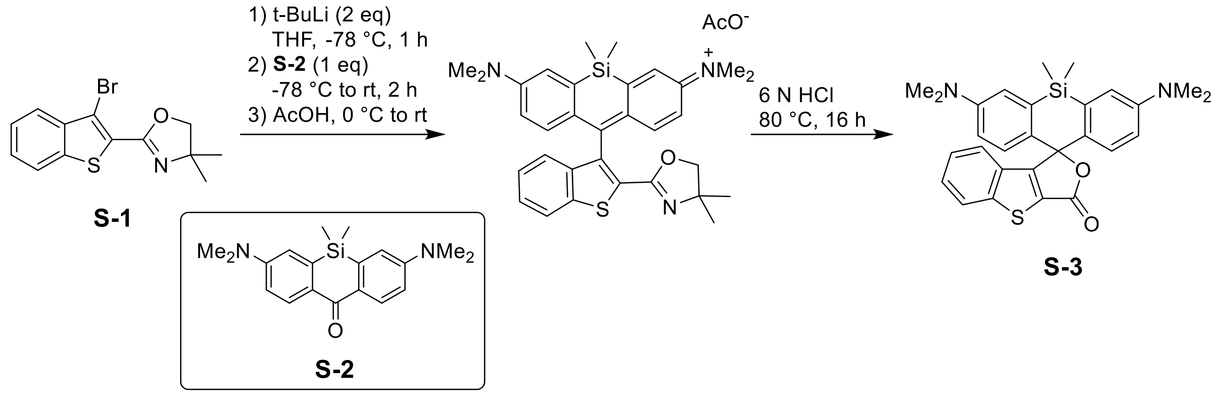

*tert*-Butyllithium (0.6 mL of 1.7 M solution in pentane, ~1 mmol, 2 equiv) was added dropwise to a stirred solution of **S-1** (310 mg, 1 mmol, 2 equiv) in anhydrous degassed THF (8 mL), cooled in dry ice-acetone bath under argon. The reaction mixture was stirred at -78 °C for 1 h, and the solution of ketone **S-2** (prepared according to the literature procedure: compound SI-7 in [4]; 162 mg, 0.5 mmol, 1 equiv) in anhydrous THF (8 mL) was added dropwise. The reaction mixture was warmed up to rt and left stirring for 2 h (light orange clear solution). It was then cooled in ice-water bath and quenched with acetic acid (2.5 mL), the intense blue-green mixture was evaporated and the residue of crude Si-pyronine intermediate was dissolved in 20 mL of 6 N HCl. The resulting orange-brown solution was stirred at 80 °C (bath temperature) overnight (16 h), cooled to rt and poured into cold 1 N NaOH (100-120 mL, adjusting the pH to ≥8). The blue-green dye was extracted with $CH_2Cl_2$ (3 × 50 mL), the combined extracts were washed with brine, dried over $Na_2SO_4$, filtered and evaporated on Celite. The product with absorption $\lambda_{max}$~650 nm was isolated by flash column chromatography (25 g Interchim SiHP 30 µm cartridge, gradient 0% to 100% A/B, A = $CH_2Cl_2$:ethanol:water 60:35:5, B = $CH_2Cl_2$) and freeze-dried from aqueous 1,4-dioxane to yield 150 mg (62%) of **S-3** as fluffy turquoise solid.

$^1H$ NMR (400 MHz, $CDCl_3$): δ 8.02 (dt, *J* = 8.3, 0.9 Hz, 1H), 7.52 (ddd, *J* = 8.3, 7.0, 1.3 Hz, 1H), 7.44 (dt, *J* = 8.0, 1.1 Hz, 1H), 7.35 (ddd, *J* = 8.1, 7.0, 1.0 Hz, 1H), 7.04 (d, *J* = 2.9 Hz, 2H), 6.69 (d, *J* = 8.8 Hz, 2H), 6.41 (dd, *J* = 8.8, 2.9 Hz, 2H), 2.96 (s, 12H), 0.67 (s, 3H), 0.65 (s, 3H).

$^{13}C$ NMR (101 MHz, $CDCl_3$): δ 165.1, 157.6, 150.1, 147.5, 139.9, 133.8, 133.2, 130.7, 129.2, 127.6, 125.8, 125.4, 124.5, 117.5, 112.7, 40.4, 1.0, -2.8.

HRMS ($C_{28}H_{28}N_2O_2SSi$): *m/z* (positive mode) = 485.1709 (found $[M+H]^+$), 485.1714 (calc.).

#### Compound S-4

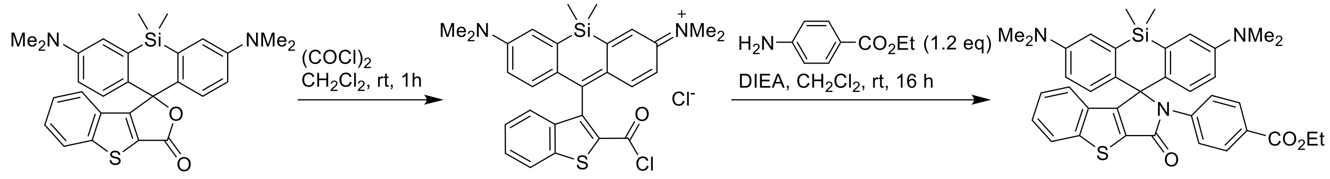

S-3 S-4

Oxalyl chloride (100 µL) was added to a stirred mixture of **S-3** (24 mg, 50 µmol) in dry $CH_2Cl_2$ (1.5 mL) and stirred at rt for 1 h. The resulting blue solution was evaporated to dryness, chased with dry $CH_2Cl_2$ (1 mL), and the residue was redissolved in dry $CH_2Cl_2$ (1.5 mL). Ethyl 4-aminobenzoate (10 mg, 60 µmol, 1.2 equiv) followed by DIEA (150 µL) were added, and the reaction mixture was left stirring overnight (16 h) at rt. It was then extracted with $CH_2Cl_2$ (3 × 20 mL) from sat. aq. $NaHCO_3$ – water mixture (1:1), the combined extracts were dried over $Na_2SO_4$, filtered and evaporated on Celite. The product was isolated by flash column chromatography (12 g Interchim SiHP 30 µm cartridge, gradient 20% to 100% EtOAc/hexane) to give 23 mg (73%) of **S-4** (purity ~85%, remainder ethyl 4-aminobenzoate), which was used in the next step without additional purification.

$^1$H NMR (400 MHz, acetone-$d_6$): δ 8.06 (dt, $J$ = 8.2, 0.9 Hz, 1H), 7.76 – 7.71 (m, 3H), 7.55 – 7.49 (m, 2H), 7.39 (ddd, $J$ = 8.2, 6.3, 2.1 Hz, 1H), 7.28 – 7.21 (m, 2H), 7.03 (d, $J$ = 2.9 Hz, 2H), 6.95 (d, $J$ = 9.0 Hz, 2H), 6.63 (dd, $J$ = 9.1, 2.9 Hz, 2H), 4.23 (q, $J$ = 7.1 Hz, 2H), 2.90 (s, 12H), 1.27 (t, J = 7.1 Hz, 3H), 0.78 (s, 3H), 0.50 (s, 3H).
$^{13}$C NMR (101 MHz, acetone-$d_6$): δ 166.1, 166.0, 160.2, 149.9, 147.4, 143.3, 135.1, 132.0, 131.94, 131.87, 130.1, 129.8, 129.0, 128.0, 127.0, 126.1, 125.4, 123.6, 122.9, 116.5, 116.0, 113.8, 74.3, 61.2, 40.1, 14.5, 0.4, -0.8.
HRMS ($C_{37}H_{37}N_3O_3SSi$): $m/z$ (positive mode) = 632.2395 (found $[M+H]^+$), 632.2398 (calc.).

#### Dye 1

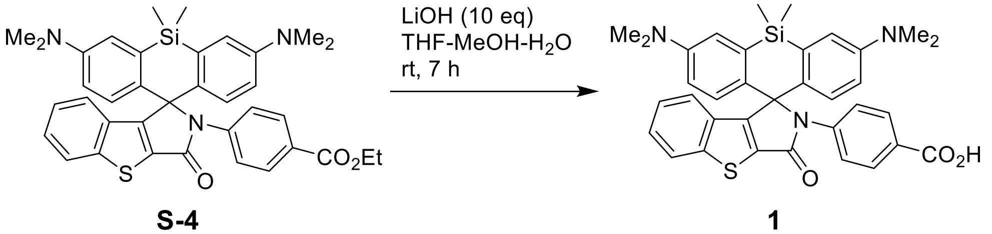

S-4 1

A solution of lithium hydroxide monohydrate (15 mg, 360 µmol) in water (300 µL) was added to the solution of **S-4** (23 mg of the crude material from the previous step) in THF (500 µL) and methanol (100 µL), and the reaction mixture was vigorously stirred at rt for 7 h. It was then quenched by addition of acetic acid (150 µL), evaporated to dryness, and the product was isolated by preparative HPLC (column: Interchim 250×21.2 mm 5 µm Uptisphere

Strategy PhC4; gradient 40/60 → 80/20 A:B, A = 0.1% v/v TFA in acetonitrile, B = 0.1% v/v TFA in water; detection at 310 and 660 nm). Fractions containing the product were evaporated (bath temperature 40 °C), and the residue was freeze-dried from aq. dioxane to give **1** as turquoise solid (25 mg, 83% over 2 steps).

$^1$H NMR (400 MHz, pyridine-$d_5$): δ 8.27 – 8.21 (m, 2H), 8.03 (dt, $J$ = 8.2, 0.9 Hz, 1H), 7.93 – 7.88 (m, 2H), 7.60 (dt, $J$ = 7.9, 1.1 Hz, 1H), 7.31 (d, $J$ = 9.0 Hz, 2H), 7.30 – 7.25 (m, 1H), 7.17 (ddd, $J$ = 8.2, 7.2, 1.1 Hz, 1H), 7.08 (d, $J$ = 2.9 Hz, 2H), 2.74 (s, 12H), 0.89 (s, 3H), 0.60 (s, 3H).

$^{13}$C NMR (101 MHz, pyridine-$d_5$): δ 168.8, 166.4, 160.4, 149.7, 147.6, 143.0, 135.4, 132.9, 132.2, 131.0, 130.0, 129.43, 129.36, 127.9, 126.2, 125.5, 124.2, 116.4, 116.2, 74.7, 40.1, 0.9, -0.3.

HRMS ($C_{35}H_{33}N_3O_3SSi$): $m/z$ (positive mode) = 604.2080 (found $[M+H]^+$), 604.2085 (calc.).

#### 1-NHS

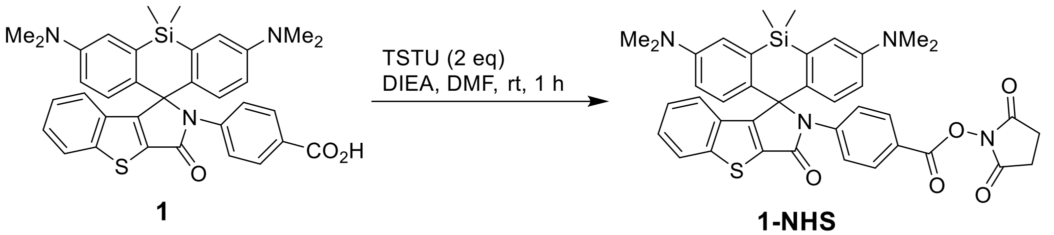

**1** → **1-NHS**

A solution of $N$,$N$,$N'$,$N'$-tetramethyl-$O$-($N$-succinimidyl)uronium tetrafluoroborate (TSTU; 22 mg, 73 µmol, 2 equiv) in DMF (200 µL) was added to the solution of **1** (22 mg, 36 µmol) and DIEA (50 µL) in DMF (200 µL), and the reaction mixture was stirred at rt for 1 h. The solvents were then evaporated *in vacuo*, the residue was redissolved in $CH_2Cl_2$, evaporated on Celite, and the product was isolated by flash column chromatography (12 g Interchim SiHP 30 µm cartridge, gradient 30% to 100% EtOAc/hexane) and freeze-dried from 1,4-dioxane to give 14.5 mg (57%) of **1-NHS** as greenish-yellow fluffy solid.

$^1$H NMR (400 MHz, acetone-$d_6$): δ 8.07 (dt, $J$ = 8.3, 0.9 Hz, 1H), 7.90 – 7.84 (m, 2H), 7.79 – 7.73 (m, 2H), 7.41 (ddd, $J$ = 8.3, 6.7, 1.7 Hz, 1H), 7.32 – 7.23 (m, 2H), 7.04 (d, $J$ = 2.9 Hz, 2H), 6.95 (d, $J$ = 9.1 Hz, 2H), 6.63 (dd, $J$ = 9.1, 2.9 Hz, 2H), 2.91 (s, 12H), 2.90 (s, 4H), 0.81 (s, 3H), 0.57 (s, 3H).

$^{13}$C NMR (101 MHz, acetone-$d_6$): δ 131.2, 128.9, 128.3, 126.2, 125.4, 123.7, 122.4, 116.7, 116.1, 40.1, 26.3, 0.4, -0.8 (indirect detection from a gHSQC experiment, only H-coupled $^{13}$C nuclei are detected).

HRMS ($C_{39}H_{36}N_4O_5SSi$): $m/z$ (positive mode) = 701.2243 (found $[M+H]^+$), 701.2248 (calc.).

#### 1-Halo

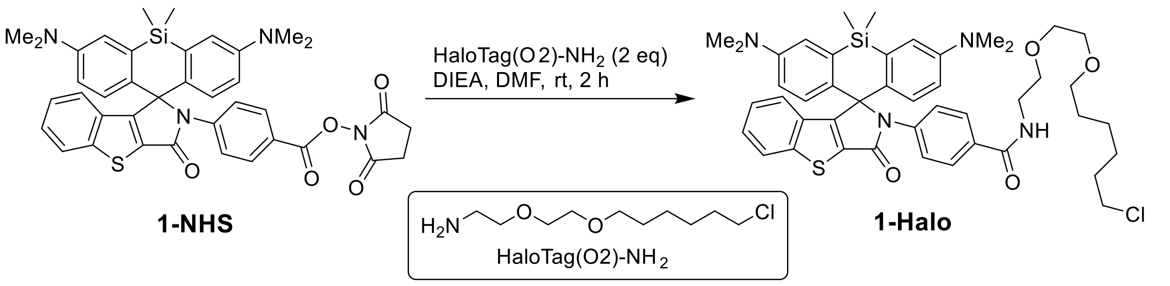

HaloTag(O2)-$NH_2$ (prepared according to the literature procedure: compound A4 in [5]; 4.5 mg, 20 µmol, 2 equiv) in DMF (50 µL) was added to the solution of **1-NHS** (7 mg, 10 µmol) and DIEA (30 µL) in DMF (100 µL), and the reaction mixture was stirred at rt for 2 h. The solvents were then evaporated *in vacuo*, and the product was isolated by preparative HPLC (column: Interchim 250×21.2 mm 5 µm Uptisphere Strategy PhC4; gradient 40/60 → 90/10 A:B, A = 0.1% v/v TFA in acetonitrile, B = 0.1% v/v TFA in water; detection at 220 and 660 nm). Fractions containing the product were evaporated (bath temperature 40 °C), and the residue was freeze-dried from dioxane to give **1-Halo** as green solid (7.5 mg, 93%).

HRMS ($C_{45}H_{53}ClN_4O_4SSi$): *m/z* (positive mode) = 809.3316 (found $[M+H]^+$), 809.3318 (calc.).

#### Compound S-5

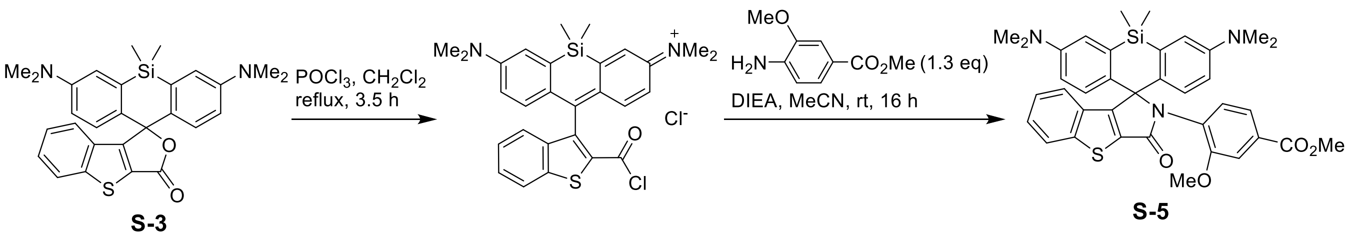

Phosphorus(V) oxychloride (0.19 mL, 2 mmol, 20 equiv) was added to a stirred mixture of **S-3** (48 mg, 0.1 mmol) in dry $CH_2Cl_2$ (5 mL), and the mixture was refluxed for 3.5 h (bath temperature 50-60 °C). The resulting blue solution was evaporated to dryness, methyl 4-amino-3-methoxybenzoate (23 mg, 0.13 mmol, 1.3 equiv), dry acetonitrile (2 mL) and DIEA (260 µL, 1.5 mmol, 15 equiv) were added to the residue, and the reaction mixture was stirred at rt overnight (16 h). The crude reaction mixture was evaporated on Celite, and the product was isolated by flash column chromatography (12 g Interchim SiHP 30 µm cartridge, gradient 20% to 100% EtOAc/hexane) to give 43 mg (66%) of **S-5** as yellowish solid.

$^1H$ NMR (400 MHz, acetone-$d_6$): δ 8.17 – 8.12 (m, 1H), 7.43 (ddd, *J* = 8.4, 7.1, 1.2 Hz, 1H), 7.38 (d, *J* = 1.8 Hz, 1H), 7.26 – 7.19 (m, 2H), 7.05 (dt, *J* = 8.0, 1.0 Hz, 1H), 6.95 (br.d, J = 9.1 Hz, 2H), 6.93 – 6.80 (br.s, 2H), 6.69 (br.dd, J = 9.1, 2.9 Hz, 2H), 6.19 (d, J = 8.1 Hz, 1H), 3.81 (s, 3H), 3.46 (s, 3H), 2.93 (s, 12H), 0.52 (s, 3H), -0.11 (s, 3H).

$^{13}C$ NMR (101 MHz, acetone-$d_6$): δ 166.6, 163.5, 159.2, 157.6, 150.1, 147.1, 136.6, 134.7, 132.8, 131.4, 131.3, 130.6, 127.5, 126.1, 125.3, 124.2, 121.5, 115.9, 115.3, 113.2, 74.6, 55.9, 52.4, 40.2, -0.40, -0.43.
HRMS ($C_{37}H_{37}N_3O_4SSi$): *m/z* (positive mode) = 648.2339 (found $[M+H]^+$), 632.2347 (calc.).

#### Dye 2

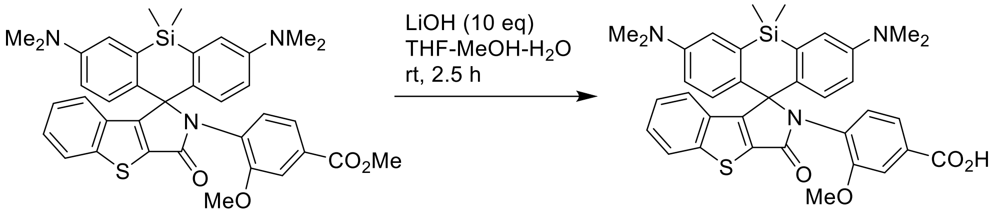

**S-5** **2**

A solution of lithium hydroxyde monohydrate (19 mg, 460 µmol) in water (300 µL) was added to the solution of **S-5** (30 mg, 46 µmol) in THF (700 µL) and methanol (150 µL), and the reaction mixture was vigorously stirred at rt for 2.5 h. It was then quenched by addition of acetic acid (400 µL), evaporated to dryness, and the product was isolated by preparative HPLC (column: Interchim 250×21.2 mm 5 µm Uptisphere Strategy PhC4; gradient 40/60 → 80/20 A:B, A = 0.1% v/v TFA in acetonitrile, B = 0.1% v/v TFA in water; detection at 220 and 670 nm). Fractions containing the product were evaporated (bath temperature 40 °C), and the residue was freeze-dried from dioxane to give **2** as dark green solid (40 mg, quant.; TFA salt, remainder dioxane).

$^1H$ NMR (400 MHz, pyridine-$d_5$): δ 8.13 (d, *J* = 8.3 Hz, 1H), 7.81 (d, *J* = 1.7 Hz, 1H), 7.74 (dd, *J* = 8.1, 1.7 Hz, 1H), 7.45 – 7.41 (m, 1H), 7.34 (ddd, *J* = 8.4, 7.1, 1.3 Hz, 1H), 7.31 (br.s, 2H), 7.11 (td, *J* = 7.6, 7.2, 1.0 Hz, 1H), 6.99 (s, 2H), 6.63 (d, *J* = 8.1 Hz, 1H), 6.59 (br.d, *J* = 8.7 Hz, 2H), 3.41 (s, 3H), 2.80 (s, 12H), 0.65 (s, 3H), 0.13 (s, 3H).
$^{13}C$ NMR (101 MHz, pyridine-$d_5$): δ 168.9, 164.4, 159.4, 157.9, 150.0, 149.8, 147.3, 136.6, 135.8, 135.6, 133.9, 133.1, 131.4, 130.6, 130.3, 127.6, 126.1, 125.5, 124.6, 124.5, 123.5, 122.4, 115.9, 115.4, 114.2, 75.0, 56.1, 40.3, 0.14, 0.06.
HRMS ($C_{36}H_{35}N_3O_4SSi$): *m/z* (positive mode) = 634.2189 (found $[M+H]^+$), 634.2190 (calc.).

#### 2-sulfoNHS

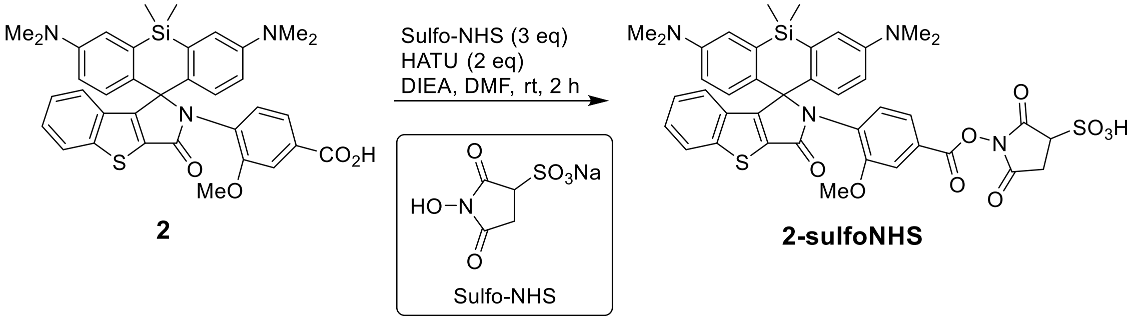

A suspension of *N*-hydroxysulfosuccinimide sodium salt (Sulfo-NHS; 6.5 mg in 50 µL of dry DMF, 30 µmol, 3 equiv) was added to the solution of **2** (7 mg, 10 µmol) and DIEA (20 µL) in DMF (300 µL) followed by addition of 1-[bis(dimethylamino)methylene]-1H-1,2,3-triazolo[4,5-b]pyridinium 3-oxid hexafluorophosphate (HATU; 7.6 mg, 20 µmol, 2 equiv) in DMF (50 µL), and the reaction mixture was stirred at rt for 2 h. The solvents were then evaporated *in vacuo*, and the product was isolated by preparative HPLC (column: Interchim 250×21.2 mm 5 µm Uptisphere Strategy PhC4; gradient 40/60 → 80/20 A:B, A = 0.1% v/v $HCO_2H$ in acetonitrile, B = 0.1% v/v $HCO_2H$ in water; detection at 220 and 670 nm). Fractions containing the product were evaporated (bath temperature 30 °C), and the residue was freeze-dried from aq. dioxane to give **2-sulfoNHS** as green solid (9 mg, quant.; remainder dioxane).

HRMS ($C_{35}H_{34}N_4O_8S_2Si$): *m/z* (positive mode) = 731.1658 (found $[M+H]^+$), 731.1660 (calc.).

#### 2-Halo

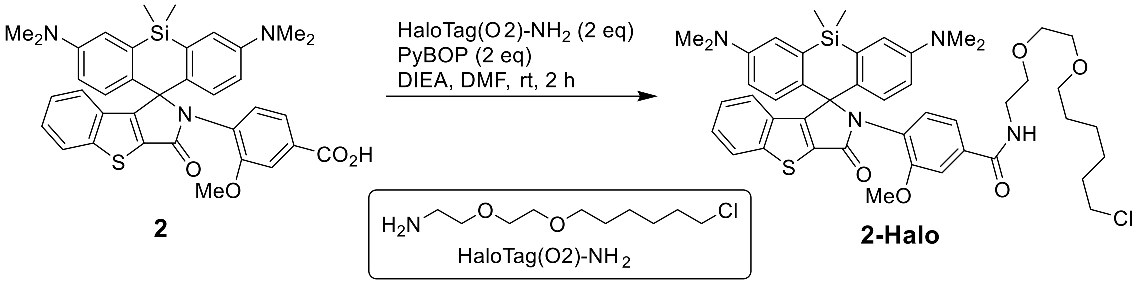

(Benzotriazol-1-yloxy)tripyrrolidinophosphonium hexafluorophosphate (PyBOP; 13 mg, 25 µmol, 2 equiv) in DMF (100 µL) was added to the solution of **2** (8 mg, 12.6 µmol), HaloTag(O2)-$NH_2$ (prepared according to the literature procedure: compound A4 in [5]; 5.7 mg, 25 µmol, 2 equiv) and DIEA (50 µL) in DMF (150 µL), and the reaction mixture was stirred at rt for 2 h. The solvents were then evaporated *in vacuo*, and the product was isolated by preparative HPLC (column: Interchim 250×21.2 mm 5 µm Uptisphere Strategy PhC4; gradient 50/50 → 100/0 A:B, A = 0.1% v/v TFA in acetonitrile, B = 0.1% v/v TFA in water; detection at 220 and 670 nm). Fractions containing the product were evaporated (bath temperature 30 °C), and the residue was freeze-dried from dioxane to give **2-Halo** as green solid (7 mg, 66%).

HRMS ($C_{46}H_{55}ClN_4O_5SSi$): *m/z* (positive mode) = 839.3422 (found $[M+H]^+$), 839.3424 (calc.).

#### Compound S-6

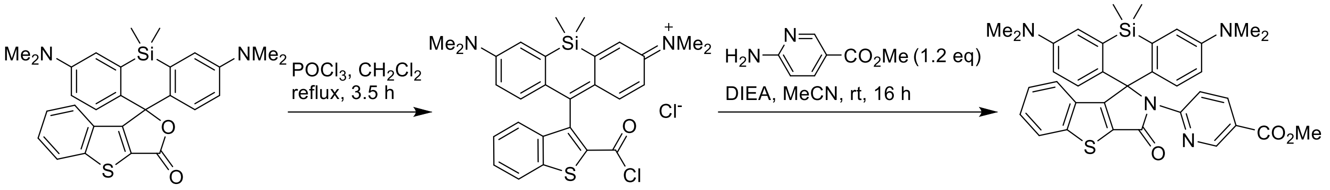

S-3

S-6

Phosphorus(V) oxychloride (0.15 mL, 1.6 mmol, 20 equiv) was added to a stirred mixture of **S-3** (39 mg, 0.08 mmol) in dry $CH_2Cl_2$ (4 mL), and the mixture was refluxed for 3.5 h (bath temperature 50-60 °C). The resulting blue-green solution was evaporated to dryness, methyl 6-aminonicotinate (15 mg, 0.1 mmol, 1.2 equiv), dry acetonitrile (1.6 mL) and DIEA (210 µL, 1.2 mmol, 15 equiv) were added to the residue, and the reaction mixture was stirred at rt overnight (16 h). The crude reaction mixture was diluted with $CH_2Cl_2$ and evaporated on Celite, and the product was isolated by flash column chromatography (12 g Interchim SiHP 30 µm cartridge, gradient 10% to 100% EtOAc/hexane) to give 36 mg (73%) of **S-6** as light yellow solid.

$^1$H NMR (400 MHz, $CDCl_3$): δ 8.63 (dd, *J* = 8.9, 0.8 Hz, 1H), 8.49 (dd, *J* = 2.4, 0.8 Hz, 1H), 8.09 (dd, *J* = 8.9, 2.4 Hz, 1H), 7.80 (dt, *J* = 8.2, 0.9 Hz, 1H), 7.29 – 7.24 (m, 1H), 7.18 (ddd, *J* = 8.1, 1.4, 0.8 Hz, 1H), 7.08 (ddd, *J* = 8.1, 7.0, 1.0 Hz, 1H), 6.88 (d, *J* = 2.9 Hz, 2H), 6.82 (d, *J* = 9.0 Hz, 2H), 6.44 (dd, *J* = 9.0, 2.9 Hz, 2H), 3.79 (s, 3H), 2.88 (s, 12H), 0.74 (s, 3H), 0.66 (s, 3H).

$^{13}$C NMR (101 MHz, $CDCl_3$): δ 166.6, 166.0, 160.4, 153.2, 149.1, 148.3, 147.3, 138.2, 134.7, 131.1, 130.9, 130.3, 127.11, 127.08, 125.2, 124.1, 123.5, 120.4, 115.7, 114.5, 113.4, 72.9, 52.0, 40.3, 0.5, -0.5.

HRMS ($C_{35}H_{34}N_4O_3SSi$): *m/z* (positive mode) = 619.2189 (found $[M+H]^+$), 619.2194 (calc.).

#### Dye 3

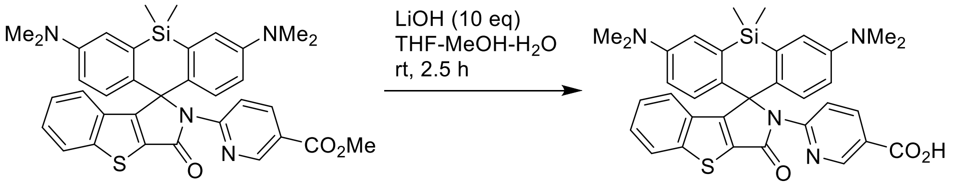

S-6

3

A solution of lithium hydroxyde monohydrate (20 mg, 480 µmol) in water (300 µL) was added to the solution of **S-6** (30 mg, 48 µmol) in THF (700 µL) and methanol (150 µL), and the reaction mixture was vigorously stirred at rt for 2.5 h. It was then quenched by addition of acetic acid (400 µL), evaporated to dryness, and the product was isolated by preparative HPLC (column: Interchim 250×21.2 mm 5 µm Uptisphere Strategy PhC4; gradient 35/65 → 75/25 A:B, A = 0.1% v/v TFA in acetonitrile, B = 0.1% v/v TFA in water; detection at 220 and 670 nm). Fractions containing the product were evaporated (bath temperature 40 °C), and the residue was freeze-dried from dioxane to give **3** as green solid (40 mg, quant.; TFA salt, remainder dioxane).

$^{1}$H NMR (400 MHz, pyridine-$d_5$): δ 9.19 (dd, $J$ = 8.9, 0.8 Hz, 1H), 8.98 (dd, $J$ = 2.3, 0.8 Hz, 1H), 8.48 (dd, $J$ = 8.9, 2.3 Hz, 1H), 7.97 (dt, $J$ = 8.2, 0.9 Hz, 1H), 7.63 (dt, $J$ = 8.0, 1.1 Hz, 1H), 7.27 – 7.21 (m, 3H), 7.19 (d, $J$ = 2.9 Hz, 2H), 7.13 (ddd, $J$ = 8.1, 7.1, 1.0 Hz, 1H), 6.46 (dd, $J$ = 9.0, 2.9 Hz, 2H), 2.70 (s, 12H), 0.96 (s, 3H), 0.90 (s, 3H).

$^{13}$C NMR (101 MHz, pyridine-$d_5$): δ 168.0, 167.2, 162.8 (q, J = 34.6 Hz, $CF_3\underline{C}O_2^-$), 161.6, 154.0, 149.3, 148.0, 139.4, 135.7, 132.1, 131.8, 131.4, 128.3, 127.9, 126.3, 125.4, 123.5, 123.4, 118.6 (q, J = 293.2 Hz, $\underline{C}F_3CO_2^-$), 116.7, 115.6, 114.3, 73.9, 40.3, 1.1, -0.1.

HRMS ($C_{34}H_{32}N_4O_3SSi$): $m/z$ (positive mode) = 605.2035 (found $[M+H]^+$), 605.2037 (calc.).

#### 3-sulfoNHS

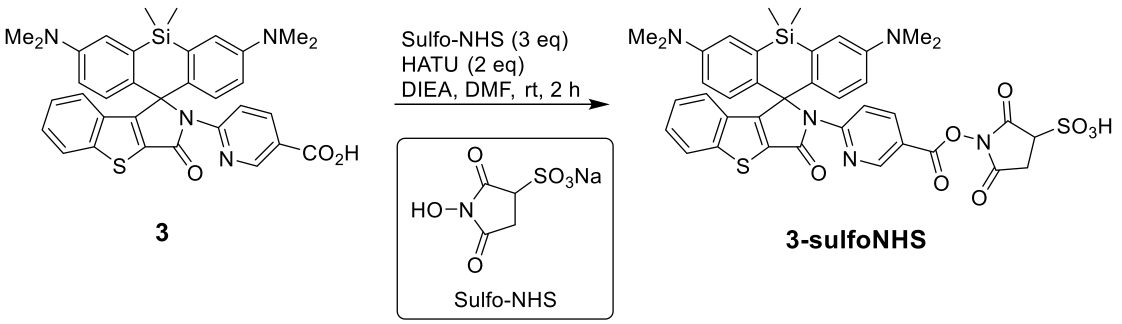

A suspension of *N*-hydroxysulfosuccinimide sodium salt (Sulfo-NHS; 6.5 mg in 50 µL of dry DMF, 30 µmol, 3 equiv) was added to the solution of **3** (7 mg, 10 µmol) and DIEA (20 µL) in DMF (300 µL) followed by addition of 1-[bis(dimethylamino)methylene]-1*H*-1,2,3-triazolo[4,5-*b*]pyridinium 3-oxid hexafluorophosphate (HATU; 7.6 mg, 20 µmol, 2 equiv) in DMF (50 µL), and the reaction mixture was stirred at rt for 2 h. The solvents were then evaporated *in vacuo*, and the product was isolated by preparative HPLC (column: Interchim 250×21.2 mm 5 µm Uptisphere Strategy PhC4; gradient 40/60 → 80/20 A:B, A = 0.1% v/v $HCO_2H$ in acetonitrile, B = 0.1% v/v $HCO_2H$ in water; detection at 220 and 670 nm). Fractions containing the product were evaporated (bath temperature 30 °C), and the residue was freeze-dried from aq. dioxane to give **3-sulfoNHS** as green solid (4.7 mg, 60%).

HRMS ($C_{38}H_{35}N_5O_8S_2Si$): $m/z$ (positive mode) = 782.1766 (found $[M+H]^+$), 782.1769 (calc.).

#### 3-Halo

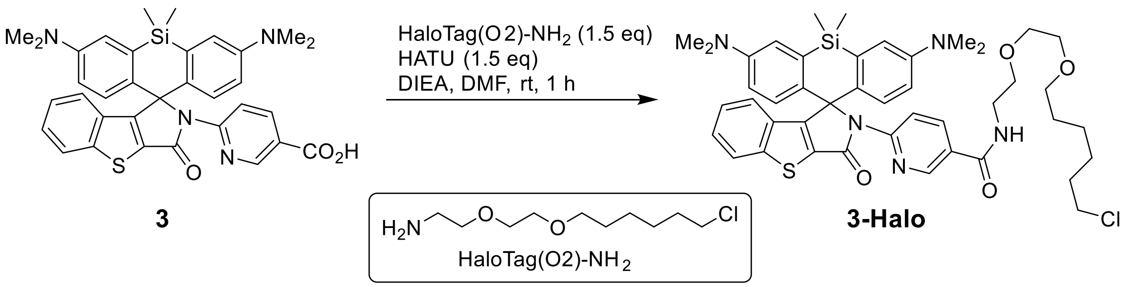

1-[bis(dimethylamino)methylene]-1*H*-1,2,3-triazolo[4,5-*b*]pyridinium 3-oxid hexafluorophosphate (HATU; 9.4 mg, 24.8 µmol, 1.5 equiv) in DMF (100 µL) was added to the solution of **3** (10 mg, 16.6 µmol), HaloTag(O2)-$NH_2$ (prepared according to the literature procedure: compound A4 in [5]; 5.6 mg, 24.8 µmol, 1.5 equiv) and DIEA (100 µL) in DMF (200 µL), and the reaction mixture was stirred at rt for 2 h. The solvents were then evaporated *in vacuo*, and the product was isolated by preparative HPLC (column: Interchim 250×21.2 mm 5 µm Uptisphere Strategy PhC4; gradient 50/50 → 100/0 A:B, A = 0.1% v/v $HCO_2H$ in acetonitrile, B = 0.1% v/v $HCO_2H$ in water; detection at 220 and 670 nm). Fractions containing the product were evaporated (bath temperature 30 °C), and the residue was freeze-dried from dioxane to give **3-Halo** as light yellow solid (8.5 mg, 63%).

HRMS ($C_{44}H_{52}ClN_5O_4SSi$): *m/z* (positive mode) = 810.3266 (found $[M+H]^+$), 810.3271 (calc.).

#### Compound S-8

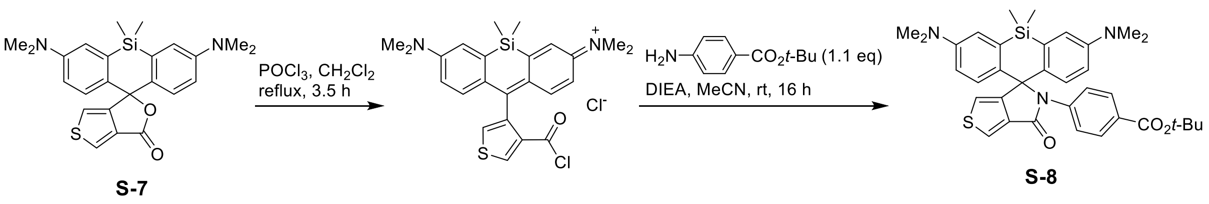

Phosphorus(V) oxychloride (0.15 mL, 1.6 mmol, 20 equiv) was added to a stirred mixture of **S-7** (prepared according to the literature procedure: compound 3c in [6]; 35 mg, 0.08 mmol) in dry $CH_2Cl_2$ (5 mL), and the mixture was refluxed for 3.5 h (bath temperature 50-60 °C). The resulting blue solution was evaporated to dryness, *tert*-butyl 4-aminobenzoate (17 mg, 0.09 mmol, 1.1 equiv), dry acetonitrile (1.5 mL) and DIEA (200 µL, 1.2 mmol, 15 equiv) were added to the residue, and the reaction mixture was stirred at rt overnight (16 h). The crude reaction mixture was diluted with $CH_2Cl_2$ and evaporated on Celite, and the product was isolated by flash column chromatography (12 g Interchim SiHP 30 µm cartridge, gradient 20% to 100% EtOAc/hexane) and freeze-dried from 1,4-dioxane to give 49 mg (73%) of **S-8** as light blue fluffy solid (purity 93%, remainder *tert*-butyl 4-aminobenzoate), which was used in the following step without additional purification.

$^1$H NMR (400 MHz, $CDCl_3$): δ 7.88 (d, *J* = 2.4 Hz, 1H), 7.70 – 7.65 (m, 2H), 7.25 – 7.21 (m, 2H), 7.00 (d, *J* = 9.0 Hz, 2H), 6.81 (d, *J* = 2.9 Hz, 2H), 6.62 (dd, *J* = 9.0, 2.9 Hz, 2H), 6.57 (d, *J* = 2.4 Hz, 1H), 2.95 (s, 12H), 1.49 (s, 9H), 0.54 (s, 3H), 0.33 (s, 3H).

$^{13}$C NMR (101 MHz, $CDCl_3$): δ 165.5, 163.8, 155.8, 148.7, 141.8, 136.2, 133.7, 133.2, 129.7, 128.6, 128.1, 123.7, 123.5, 115.7, 115.4, 115.2, 80.7, 72.5, 40.3, 28.3, 0.6, -0.6.

HRMS ($C_{35}H_{39}N_3O_3SSi$): *m/z* (positive mode) = 610.2549 (found $[M+H]^+$), 610.2554 (calc.).

#### Dye 4

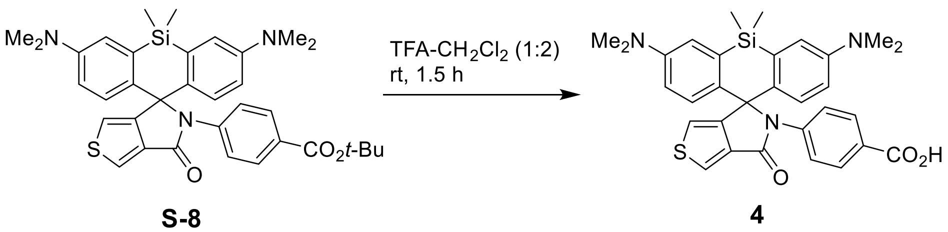

Trifluoroacetic acid (200 µL) was added to a blue solution of **S-8** (40 mg, 66 µmol) in $CH_2Cl_2$ (400 µL), and the resulting yellow-orange solution was stirred at rt for 1.5 h. The reaction mixture was diluted with $CH_2Cl_2$ – toluene (1:1, 5 mL) and evaporated; the product was isolated from the residue by preparative HPLC (column: Interchim 250×21.2 mm 5 µm Uptisphere Strategy PhC4; gradient 40/60 → 75/25 A:B, A = 0.1% v/v TFA in acetonitrile, B = 0.1% v/v TFA in water; detection at 220 and 650 nm). Fractions containing the product were evaporated (bath temperature 40 °C), and the residue was freeze-dried from aq. dioxane to give **4** as dark blue solid (48 mg, quant.; TFA salt, remainder dioxane).

$^1$H NMR (400 MHz, pyridine-*d*$_5$): δ 8.41 (d, *J* = 2.4 Hz, 1H), 8.22 – 8.17 (m, 2H), 8.04 – 7.98 (m, 2H), 7.35 (d, *J* = 9.0 Hz, 2H), 7.09 (d, *J* = 2.9 Hz, 2H), 7.02 (d, *J* = 2.4 Hz, 1H), 6.63 (dd, *J* = 9.0, 2.9 Hz, 2H), 2.77 (s, 12H), 0.71 (s, 3H), 0.64 (s, 3H).

$^{13}$C NMR (101 MHz, pyridine-*d*$_5$): δ 168.9, 164.6, 162.8 (q, J = 34.6 Hz, $CF_3\underline{C}O_2^-$), 157.2, 149.6, 143.6, 137.1, 134.4, 134.3, 130.9, 129.1, 128.7, 125.2, 118.6 (q, J = 293.2 Hz, $\underline{C}F_3CO_2^-$), 116.9, 116.8, 116.3, 73.4, 40.3, 1.1, 0.0.

HRMS ($C_{31}H_{31}N_3O_3SSi$): *m/z* (positive mode) = 554.1921 (found $[M+H]^+$), 554.1928 (calc.).

#### 4-Halo

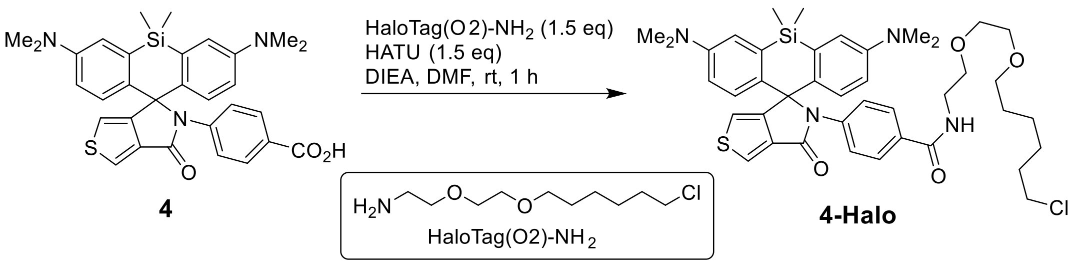

A solution of 1-[bis(dimethylamino)methylene]-1H-1,2,3-triazolo[4,5-b]pyridinium 3-oxid hexafluorophosphate (HATU; 9.4 mg, 25 µmol, 1.5 equiv) in DMF (100 µL) was added to the solution of **4** (TFA salt; 10 mg, 15 µmol), HaloTag(O2)-$NH_2$ (prepared according to the literature procedure: compound A4 in [5]; 5.6 mg, 25 µmol, 1.5 equiv) and DIEA (100 µL) in DMF (200 µL), and the reaction mixture was stirred at rt for 1 h. The solvents were then evaporated *in vacuo*, and the product was isolated by preparative HPLC (column: Interchim 250×21.2 mm 5 µm Uptisphere Strategy PhC4; gradient 50/50 → 100/0 A:B, A = 0.1% v/v $HCO_2H$ in acetonitrile, B = 0.1% v/v $HCO_2H$ in water; detection at 220 and 670 nm). Fractions containing the product were evaporated (bath temperature 40 °C), and the residue was freeze-dried from dioxane to give **4-Halo** as light yellow solid (8.5 mg, 63%).

$^1$H NMR (400 MHz, pyridine-$d_5$): δ 8.78 (t, *J* = 5.7 Hz, 1H), 8.39 (d, *J* = 2.4 Hz, 1H), 8.07 – 8.00 (m, 2H), 7.98 – 7.89 (m, 2H), 7.32 (d, *J* = 9.0 Hz, 2H), 7.08 (d, *J* = 2.9 Hz, 2H), 7.01 (d, *J* = 2.4 Hz, 1H), 6.63 (dd, *J* = 9.0, 2.9 Hz, 2H), 3.77 – 3.70 (m, 2H), 3.69 – 3.64 (m, 2H), 3.61 – 3.56 (m, 2H), 3.55 – 3.47 (m, 4H), 3.35 (t, *J* = 6.5 Hz, 2H), 2.79 (s, 12H), 1.68 – 1.57 (m, 2H), 1.48 (p, J = 6.7 Hz, 2H), 1.37 – 1.20 (m, 4H), 0.70 (s, 3H), 0.58 (s, 3H).
$^{13}$C NMR (101 MHz, pyridine-$d_5$): δ 167.0, 164.0, 156.6, 149.1, 141.7, 136.7, 133.9, 133.8, 131.5, 128.6, 128.0, 124.5, 116.4, 116.2, 115.8, 72.8, 71.1, 70.6, 70.4, 70.3, 45.5, 40.3, 39.8, 32.8, 30.8, 29.9, 26.9, 25.7, 0.6, -0.5.
HRMS ($C_{41}H_{51}ClN_4O_4SSi$): *m/z* (positive mode) = 759.3159 (found $[M+H]^+$), 759.3162 (calc.).

#### 4-NHS

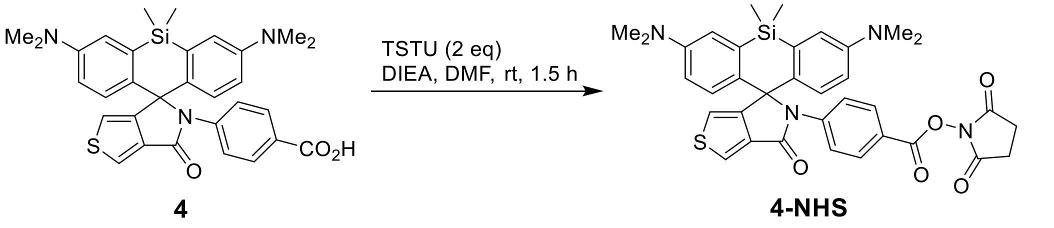

A solution of *N*,*N*,*N′*,*N′*-tetramethyl-*O*-(*N*-succinimidyl)uronium tetrafluoroborate (TSTU; 64 mg, 0.21 mmol, 2 equiv) in DMF (200 µL) was added to the solution of **4** (TFA salt; 67 mg, 0.1 mmol) and DIEA (300 µL) in DMF (300 µL), and the reaction mixture was stirred at rt for 1.5 h. The solvents were then evaporated *in vacuo*, and the product was isolated by flash column chromatography (12 g Interchim SiHP 30 µm cartridge, gradient 20% to 100%

EtOAc/hexane) and freeze-dried from 1,4-dioxane to give 60 mg (78%) of **4-NHS** as light green fluffy solid (purity 85%), which was used in the following step without additional purification.

$^{1}$H NMR (400 MHz, $CD_3CN$): δ 7.99 (d, *J* = 2.4 Hz, 1H), 7.84 – 7.79 (m, 2H), 7.66 – 7.61 (m, 2H), 6.98 (d, *J* = 2.9 Hz, 2H), 6.94 (d, *J* = 9.0 Hz, 2H), 6.64 (dd, *J* = 9.0, 2.9 Hz, 2H), 6.61 (d, *J* = 2.4 Hz, 1H), 2.91 (s, 12H), 2.79 (s, 4H), 0.58 (s, 3H), 0.55 (s, 3H).

$^{13}$C NMR (101 MHz, $CD_3CN$): δ 171.1, 164.6, 162.4, 156.6, 150.0, 145.8, 135.8, 134.4, 133.5, 131.2, 128.4, 125.9, 123.3, 120.7, 117.2, 116.5, 116.1, 73.3, 40.4, 26.4, -0.8.

HRMS ($C_{35}H_{34}N_4O_5SSi$): *m/z* (positive mode) = 651.2085 (found $[M+H]^+$), 651.2092 (calc.).

#### 4-Maleimide

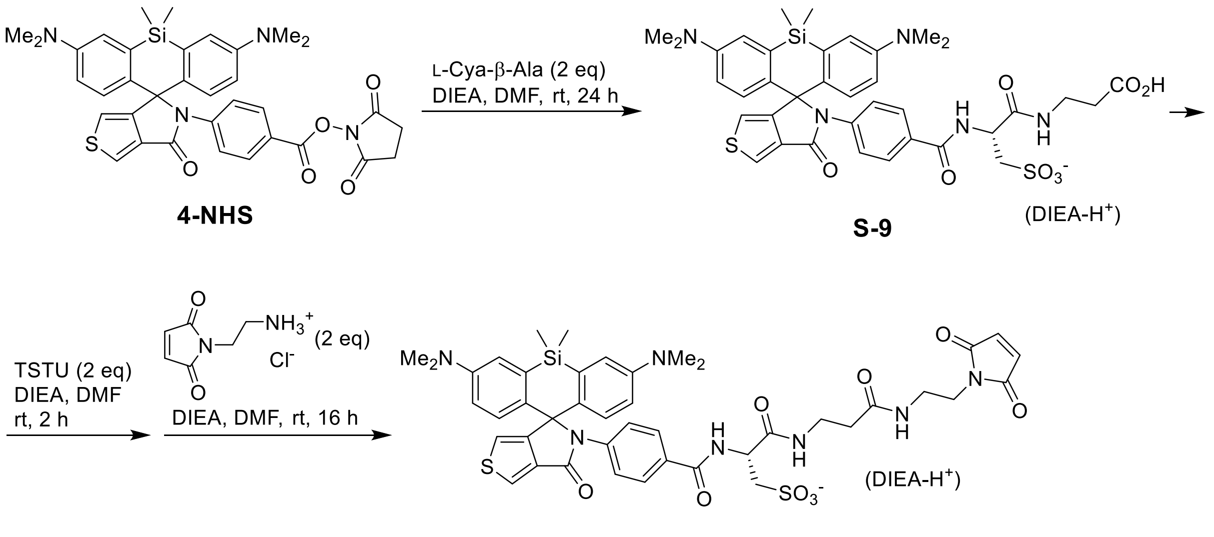

**4-Maleimide**

**4-NHS** (40 mg, 61 µmol) and L-Cya-β-Ala (prepared according to the literature procedure: compound 3 in [7]; 30 mg, 122 µmol, 2 equiv) were mixed in dry DMF (200 µL) and DIEA (100 µL), and the reaction mixture was vigorously stirred for 24 h. The solvents were then evaporated *in vacuo*, and the intermediate **S-9** was isolated by preparative HPLC (column: Interchim 250×21.2 mm 5 µm Uptisphere Strategy PhC4; gradient 20/80 → 70/30 A:B, A = 0.1% v/v $HCO_2H$ in acetonitrile, B = 0.1% v/v $HCO_2H$ in water; detection at 220, 290 and 660 nm). Fractions containing the product were evaporated (bath temperature 40 °C), and the residue was freeze-dried from aq. dioxane to give 11 mg (27%) of dark blue-green solid which was taken into the next step without further characterization.

One-half of the obtained **S-9** (5 mg, 6.3 µmol) was dissolved in dry DMF (120 µL) and DIEA (30 µL), and TSTU (2×30 µL of 6 mg in 100 µL DMF stock solution/1000; 12.6 µmol, 2 equiv) was added in two portions in 1 h intervals. Afterwards, 1-(2-aminoethyl)maleimide hydrochloride (2.2 mg, 12.6 µmol, 2 equiv) in dry DMF (100 µL) was added, followed by additional DIEA (30 µL), and the reaction mixture was stirred at rt overnight (16 h). The solvents were then evaporated *in vacuo*, and the product was isolated by preparative HPLC (column: Interchim 250×21.2 mm 5 µm Uptisphere Strategy PhC4; gradient 20/80 → 60/40 A:B, A = 0.1% v/v $HCO_2H$ in acetonitrile, B = 0.1% v/v $HCO_2H$ in water; detection at 220, 254 and 650 nm). Fractions containing the product were evaporated (bath temperature 40 °C), and the residue was freeze-dried from aq. dioxane to give 4.8 mg (85%) of **4-Maleimide**.

$^1$H NMR (400 MHz, pyridine-$d_5$): δ 9.71 (d, *J* = 7.1 Hz, 1H, NH), 9.03 (t, *J* = 6.0 Hz, 1H, NH), 8.77 (br.t, *J* = 6.1 Hz, 1H, NH), 8.35 (d, *J* = 2.4 Hz, 1H), 8.02 – 7.94 (m, 2H), 7.88 – 7.82 (m, 2H), 7.28 (dd, *J* = 9.0, 1.5 Hz, 2H), 7.08 (app.t, *J* = 3.2 Hz, 2H), 6.97 (d, *J* = 2.4 Hz, 1H), 6.68 (s, 2H), 6.64 (ddd, *J* = 9.0, 3.0, 0.9 Hz, 2H), 5.51 (dt, *J* = 6.9, 5.3 Hz, 1H), 4.06 (dd, *J* = 13.8, 5.3 Hz, 1H), 3.97 – 3.81 (m, 3H), 3.80 – 3.65 (m, 4H), 3.58 – 3.41 (m, 2H), 2.80 (s, 6H), 2.79 (s, 6H), 2.76 – 2.66 (m, 1H), 2.46 (ddd, *J* = 13.3, 6.7, 4.0 Hz, 1H), 0.68 (s, 3H), 0.58 (s, 3H).

$^{13}$C NMR (101 MHz, pyridine-$d_5$): δ 134.4, 128.6, 128.0, 124.4, 123.3, 116.3, 116.2, 115.8, 52.5, 52.3, 39.9, 38.6, 38.4, 37.14, 37.09, 0.6, -0.5 (indirect detection from a gHSQC experiment, only H-coupled $^{13}$C nuclei are detected).

HRMS ($C_{43}H_{47}N_7O_9S_2Si$): *m/z* (positive mode) = 898.2720 (found $[M+H]^+$), 898.2719 (calc.).

#### Compound S-10

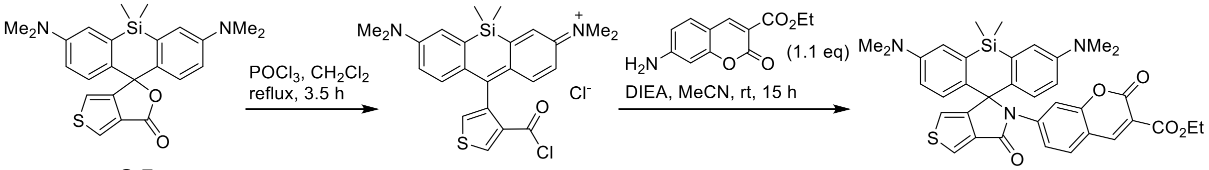

Phosphorus(V) oxychloride (0.19 mL, 2 mmol, 20 equiv) was added to a stirred mixture of **S-7** (prepared according to the literature procedure: compound 3c in [6]; 44 mg, 0.1 mmol) in dry $CH_2Cl_2$ (5 mL), and the mixture was refluxed for 3.5 h (bath temperature 50-60 °C). The resulting blue solution was evaporated to dryness, ethyl 7-aminocoumarin-3-carboxyate (prepared according to the literature procedure: compound S18 in [8]; 26 mg, 0.11 mmol, 1.1 equiv), dry acetonitrile (1.8 mL) and DIEA (260 µL, 1.5 mmol, 15 equiv) were added to the residue, and the reaction mixture was stirred at rt overnight (15 h). The crude reaction mixture was diluted with $CH_2Cl_2$, evaporated on silica gel, and the product was isolated by flash column chromatography (12 g Interchim SiHP 30 µm cartridge, gradient 25% to 100% EtOAc/hexane) and freeze-dried from 1,4-dioxane to give 35 mg (54%) of **S-10** as yellow solid.

$^1$H NMR (400 MHz, $CDCl_3$): δ 8.32 (d, *J* = 0.7 Hz, 1H), 7.92 (d, *J* = 2.4 Hz, 1H), 7.53 (dd, *J* = 8.7, 2.1 Hz, 1H), 7.29 (d, *J* = 8.7 Hz, 1H), 7.27 (d, *J* = 2.1 Hz, 1H), 6.97 (d, *J* = 9.0 Hz, 2H), 6.84 (d, *J* = 2.9 Hz, 2H), 6.61 (dd, *J* = 9.0, 2.9 Hz, 2H), 6.55 (d, *J* = 2.4 Hz, 1H), 4.35 (q, *J* = 7.1 Hz, 2H), 2.96 (s, 12H), 1.35 (t, *J* = 7.1 Hz, 3H), 0.57 (s, 3H), 0.49 (s, 3H).

$^{13}$C NMR (101 MHz, $CDCl_3$): δ 164.0, 163.4, 157.2, 155.6, 155.4, 148.8, 148.4, 144.4, 135.0, 133.5, 132.5, 129.1, 128.1, 124.6, 120.3, 116.3, 115.9, 115.5, 115.3, 114.3, 110.5, 73.0, 61.9, 40.2, 14.3, 0.9, -0.6.

HRMS ($C_{36}H_{35}N_3O_5SSi$): *m/z* (positive mode) = 650.2134 (found $[M+H]^+$), 650.2139 (calc.).

#### Dye 5

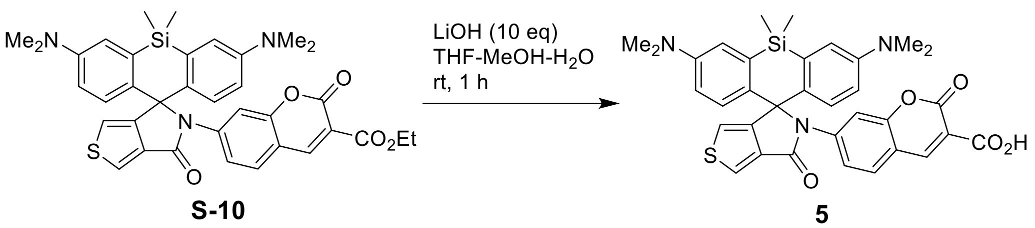

A solution of lithium hydroxide monohydrate (30 mg, 720 µmol) in water (600 µL) was added to the solution of **S-10** (30 mg, 46 µmol) in THF (2 mL) and methanol (600 µL), and the reaction mixture was vigorously stirred at rt for 1 h. It was then quenched by addition of acetic acid (1 mL), evaporated to dryness, and the product was isolated by preparative HPLC (column: Interchim 250×21.2 mm 5 µm Uptisphere Strategy PhC4; gradient 30/70 → 80/20 A:B, A = 0.1% v/v $HCO_2H$ in acetonitrile, B = 0.1% v/v $HCO_2H$ in water; detection at 220 and 660 nm). Fractions containing the product were evaporated (bath temperature 40 °C), and the residue was freeze-dried from dioxane to give **5** as green-yellow solid (28 mg, 97%).

$^1$H NMR (400 MHz, pyridine-*d*$_5$): δ 8.54 (s, 1H), 8.46 (d, *J* = 2.4 Hz, 1H), 8.12 – 8.05 (m, 2H), 7.37 (d, *J* = 8.6 Hz, 1H), 7.36 (d, *J* = 9.0 Hz, 2H), 7.14 (d, *J* = 2.9 Hz, 2H), 7.02 (d, *J* = 2.4 Hz, 1H), 6.69 (dd, *J* = 9.0, 2.9 Hz, 2H), 2.77 (s, 12H), 0.83 (s, 3H), 0.74 (s, 3H).
$^{13}$C NMR (101 MHz, pyridine-*d*$_5$): δ 166.1, 164.9, 158.4, 156.8, 156.1, 149.7, 148.2, 145.0, 134.1, 133.8, 130.1, 128.6, 126.1, 119.5, 118.5, 117.0, 116.3, 115.1, 109.9, 73.6, 40.2, 1.2, -0.2.
HRMS ($C_{34}H_{31}N_3O_5SSi$): *m/z* (positive mode) = 622.1829 (found $[M+H]^+$), 622.1826 (calc.).

#### 5-NHS

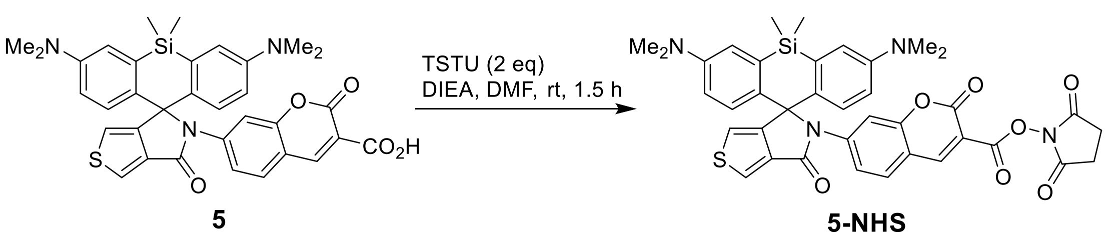

A solution of *N*,*N*,*N′*,*N′*-tetramethyl-*O*-(*N*-succinimidyl)uronium tetrafluoroborate (TSTU; 8 mg, 26 µmol, 2 equiv) in DMF (50 µL) was added to the solution of **5** (8.1 mg, 13 µmol) and DIEA (40 µL) in DMF (100 µL), and the reaction mixture was stirred at rt for 1.5 h. The solvents were then evaporated *in vacuo*, and the product was isolated by preparative HPLC (column: Interchim 250×21.2 mm 5 µm Uptisphere Strategy PhC4; gradient 40/60 → 80/20 A:B,

A = 0.1% v/v $HCO_2H$ in acetonitrile, B = 0.1% v/v $HCO_2H$ in water; detection at 220 and 660 nm). Fractions containing the product were evaporated (bath temperature 30 °C), and the residue was freeze-dried from aq. dioxane to give **5-NHS** as green solid (7 mg, 75%).

HRMS ($C_{38}H_{34}N_4O_7SSi$): *m/z* (positive mode) = 719.1989 (found $[M+H]^+$), 719.1990 (calc.).

#### 5-Halo

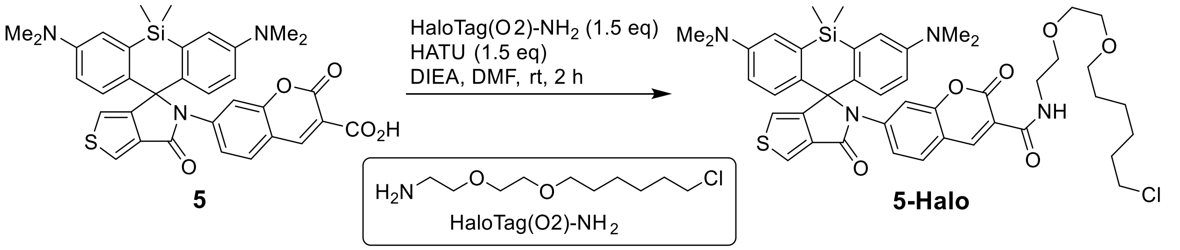

A solution of 1-[bis(dimethylamino)methylene]-1H-1,2,3-triazolo[4,5-b]pyridinium 3-oxid hexafluorophosphate (HATU; 5.7 mg, 15 µmol, 1.5 equiv) in DMF (50 µL) was added to the solution of **5** (6.2 mg, 10 µmol), HaloTag(O2)-$NH_2$ (prepared according to the literature procedure: compound A4 in [5]; 3.4 mg, 15 µmol, 1.5 equiv) and DIEA (30 µL) in DMF (200 µL), and the reaction mixture was stirred at rt for 2 h. The solvents were then evaporated *in vacuo*, and the product was isolated by preparative HPLC (column: Interchim 250×21.2 mm 5 µm Uptisphere Strategy PhC4; gradient 50/50 → 100/0 A:B, A = 0.1% v/v $HCO_2H$ in acetonitrile, B = 0.1% v/v $HCO_2H$ in water; detection at 220 and 660 nm). Fractions containing the product were evaporated (bath temperature 40 °C), and the residue was freeze-dried from dioxane to give **5-Halo** as green solid (7 mg, 85%).

HRMS ($C_{41}H_{51}ClN_4O_4SSi$): *m/z* (positive mode) = 759.3159 (found $[M+H]^+$), 759.3162 (calc.).

#### Compound S-11

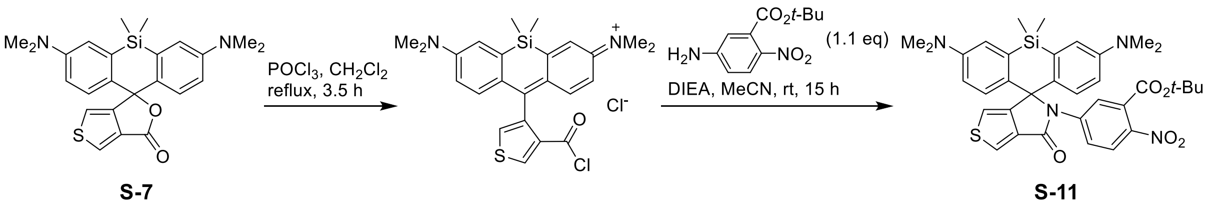

Phosphorus(V) oxychloride (0.15 mL, 1.6 mmol, 20 equiv) was added to a stirred mixture of **S-7** (prepared according to the literature procedure: compound 3c in [6]; 35 mg, 0.08 mmol) in dry $CH_2Cl_2$ (5 mL), and the mixture was refluxed for 4 h (bath temperature 50-60 °C). The resulting blue solution was evaporated to dryness, *tert*-butyl 5-amino-2-nitrobenzoate (prepared according to the literature procedure: compound 2A in [9]; (17 mg, 0.09 mmol, 1.1 equiv), dry acetonitrile (1.5 mL) and DIEA (200 µL, 1.2 mmol, 15 equiv) were added to the residue, and the reaction mixture was stirred at rt overnight (15 h). The crude reaction mixture was diluted with $CH_2Cl_2$ and evaporated on Celite, and the product was isolated by flash column chromatography (12 g Interchim SiHP 30 µm cartridge, gradient 20% to 100% EtOAc/hexane) and freeze-dried from 1,4-dioxane to give 39 mg (74%) of **S-11** as yellow solid.

$^1$H NMR (400 MHz, $CDCl_3$): δ 7.94 (d, *J* = 2.4 Hz, 1H), 7.91 (d, *J* = 2.4 Hz, 1H), 7.60 (d, *J* = 9.1 Hz, 1H), 7.31 (dd, *J* = 9.1, 2.5 Hz, 1H), 6.95 (d, *J* = 9.0 Hz, 2H), 6.85 (d, *J* = 2.9 Hz, 2H), 6.61 (dd, *J* = 9.0, 2.9 Hz, 2H), 6.57 (d, *J* = 2.4 Hz, 1H), 2.95 (s, 12H), 1.49 (s, 9H), 0.57 (s, 3H), 0.50 (s, 3H).

$^{13}$C NMR (101 MHz, $CDCl_3$): δ 164.6, 164.0, 155.3, 148.9, 142.9, 142.3, 134.9, 133.4, 132.3, 130.4, 127.9, 124.7, 124.2, 124.0, 122.8, 115.9, 115.6, 115.4, 83.3, 72.8, 40.2, 27.8, 1.1, -0.8.

HRMS ($C_{35}H_{38}N_4O_5SSi$): *m/z* (positive mode) = 655.2408 (found $[M+H]^+$), 655.2405 (calc.).

#### Dye 6

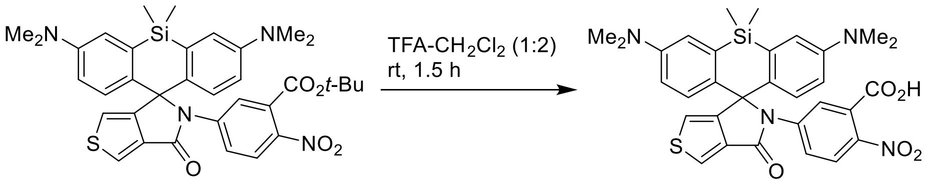

**S-11** **6**

Trifluoroacetic acid (200 µL) was added to a solution of **S-11** (35 mg, 53 µmol) in $CH_2Cl_2$ (400 µL), and the resulting orange-brown solution was stirred at rt for 1.5 h. The reaction mixture was diluted and chased twice with $CH_2Cl_2$–toluene (1:1, 6 mL); the product residue was freeze-dried from aq. dioxane to give **6** as blue solid (42 mg, quant.; TFA salt, remainder dioxane).

$^1$H NMR (400 MHz, pyridine-$d_5$): δ 9.11 (d, *J* = 2.4 Hz, 1H), 8.44 (d, *J* = 2.4 Hz, 1H), 7.94 (dd, *J* = 9.1, 2.4 Hz, 1H), 7.70 (d, *J* = 9.1 Hz, 1H), 7.31 (d, *J* = 9.0 Hz, 2H), 7.14 (d, *J* = 2.9 Hz, 2H), 7.01 (d, *J* = 2.4 Hz, 1H), 6.64 (dd, *J* = 9.0, 2.9 Hz, 2H), 2.75 (s, 12H), 0.84 (s, 3H), 0.74 (s, 3H).

$^{13}$C NMR (101 MHz, pyridine-$d_5$): δ 168.8, 164.9, 162.8 (q, J = 34.6 Hz, $CF_3\underline{C}O_2^-$), 156.6, 149.8, 144.4, 143.8, 135.7, 134.2, 133.5, 131.9, 128.3, 126.4, 125.1, 123.7, 118.6 (q, J = 293.2 Hz, $\underline{C}F_3CO_2^-$), 117.1, 117.0, 116.4, 73.6, 40.3, 1.5, -0.3.

HRMS ($C_{31}H_{30}N_4O_5SSi$): *m/z* (positive mode) = 599.1776 (found $[M+H]^+$), 599.1779 (calc.).

#### 6-NHS

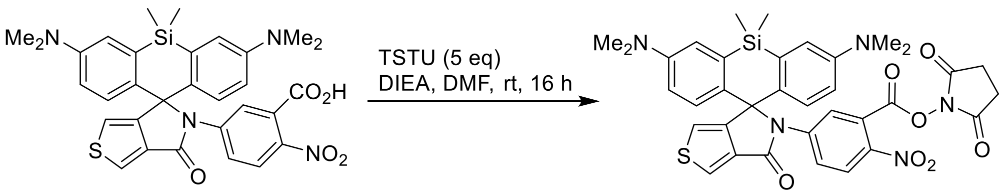

**6** **6-NHS**

*N*,*N*,*N′*,*N′*-Tetramethyl-*O*-(*N*-succinimidyl)uronium tetrafluoroborate (TSTU; 21 mg, 70 µmol, 5 equiv) was added to the solution of **6** (10 mg, 14 µmol) and DIEA (60 µL) in DMF (200 µL), and the reaction mixture was stirred at rt overnight (16 h; the conversion did not exceed 50% by LC-MS analysis and was not improved by further addition of TSTU). The solvents were then evaporated *in vacuo*, and the product was isolated by preparative HPLC (column: Interchim 250×21.2 mm 5 µm Uptisphere Strategy PhC4; gradient 40/60 → 80/20 A:B, A = 0.1% v/v TFA in acetonitrile, B = 0.1% v/v TFA in water; detection at 220 and 660 nm). Fractions containing the product were evaporated (bath temperature 30 °C), and the residue was freeze-dried from dioxane to give **6-NHS** as blue solid (4.5 mg, 46%).

HRMS ($C_{35}H_{33}N_5O_7SSi$): *m/z* (positive mode) = 696.1936 (found $[M+H]^+$), 696.1943 (calc.).

#### 6-Halo

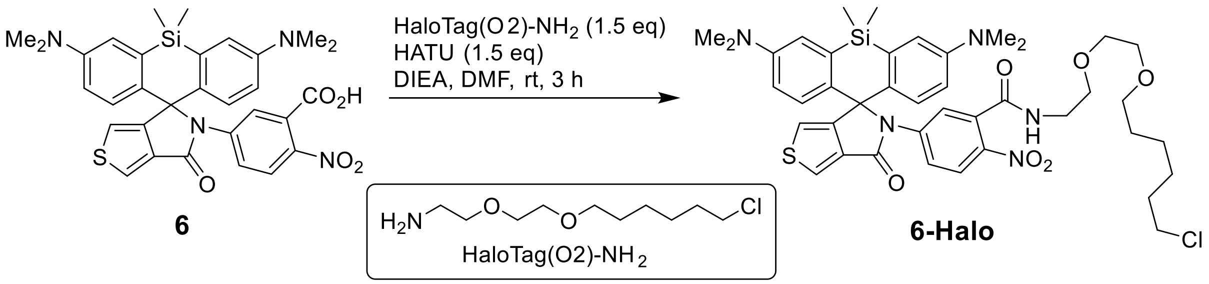

**6** **6-Halo**

A solution of 1-[bis(dimethylamino)methylene]-1H-1,2,3-triazolo[4,5-b]pyridinium 3-oxid hexafluorophosphate (HATU; 5.7 mg, 15 µmol, 1.5 equiv) in DMF (50 µL) was added to the solution of **6** (7 mg, 10 µmol), HaloTag(O2)-$NH_2$ (prepared according to the literature procedure: compound A4 in [5]; 3.4 mg, 15 µmol, 1.5 equiv) and DIEA (20 µL) in DMF (150 µL), and the reaction mixture was stirred at rt for 3 h. The solvents were then evaporated *in vacuo*, the residue was diluted with $CH_2Cl_2$ and evaporated on Celite, and the product was isolated by flash column chromatography (12 g Interchim SiHP 30 µm cartridge, gradient 0% to 100% EtOAc/$CH_2Cl_2$) and freeze-dried from 1,4-dioxane to give 7.8 mg (97%) of **6-Halo** as light-green solid.

HRMS ($C_{41}H_{50}ClN_5O_6SSi$): *m/z* (positive mode) = 804.3007 (found $[M+H]^+$), 804.3012 (calc.).
